# Cholesterol-dependent enrichment of understudied erythrocytic stages of human *Plasmodium* parasites

**DOI:** 10.1101/474338

**Authors:** Audrey C. Brown, Christopher C. Moore, Jennifer L. Guler

## Abstract

*Plasmodium* protozoan parasites undergo rounds of asexual replication inside human erythrocytes, progressing from ring stage, to trophozoites and schizonts, before egress and reinvasion. Given the discovery of ring-specific artemisinin tolerance and quiescence in *Plasmodium falciparum*, there is great urgency to better understand ring stage biology. However, the lack of an effective enrichment method has left rings and related parasite stages understudied compared to their late stage counterparts, which can be easily isolated due to their paramagnetic properties. Here, a method for separating *all Plasmodium* infected erythrocytes from uninfected erythrocytes is presented. This approach takes advantage of streptolysin-O (SLO) to preferentially lyse uninfected erythrocytes as previously shown by Jackson, *et al.* Following lytic treatment, Percoll gradient centrifugation removes lysed cells, leaving an intact cell population enriched in infected erythrocytes. This <u>SLO</u>-<u>Pe</u>rcoll (SLOPE) method is effective on stages from the entire erythrocytic cycle, including previously inaccessible forms such as circulating rings from malaria-infected patients and artemisinin-induced quiescent parasites. Furthermore, the utility of SLOPE is extended to multiple media formulations used for the propagation of two human *Plasmodium* species. The alteration of external cholesterol levels modulates SLOPE effectiveness, demonstrating the role of erythrocyte membrane cholesterol in lytic discrimination. Importantly, enrichment does not impact parasite viability, which establishes the non-toxic nature of SLOPE. Targeted metabolomics of SLOPE-enriched ring stage samples confirms the impact on treated samples; parasite-derived metabolites are increased and contaminating host material is reduced compared to non-enriched samples.

**Importance:** Malaria is caused by infection with protozoan *Plasmodium* parasites and is responsible for over 400,000 deaths annually. The availability of effective antimalarial drugs is critical to the reduction of malaria-related mortality, yet widespread resistance highlights the need for the continued study of *Plasmodium* biology. The SLOPE method is an accessible, scalable, rapid (30-40min), and non-toxic enrichment method that is broadly effective on many erythrocytic stages. This method is ideal for use upstream of a variety of sensitive analyses, which will increase experimental quality in virtually all areas of asexual *Plasmodium* parasite research. Further, because the consumption of cholesterol is a common characteristic of other intracellular parasites (both bacteria and other protozoa), SLOPE holds potential for extension to other relevant pathogens.

## Introduction

Malaria, caused by protozoan parasites of the *Plasmodium* genus, is a continuing threat to global health. A total of five *Plasmodium* species cause malaria in humans, with *Plasmodium falciparum* being responsible for the large majority of malaria morbidity and mortality (1). While the global malaria burden has decreased over the past decade, the emergence and spread of antimalarial resistant *Plasmodium* threatens to undo this progress and emphasizes the dire need to understand more about the biology of this parasite. The current World Health Organization recommendation for treatment of malaria is artemisinin combination therapy (2). However, clinical resistance has now been reported to both artemisinin and almost all of its partner drugs (3–5).

All symptoms of malaria, including cyclical fevers and hypoglycemia, occur due to the asexual replication cycle of the parasite within human erythrocytes (Fig. 1A). Parasites undergo rounds of replication progressing from the ring stage, to trophozoites and schizonts, before rupturing from host erythrocytes to release merozoites, which go on to invade new erythrocytes and continue the cycle of infection (6). Many studies aiming to understand the biology of asexual *Plasmodium* are performed only on late stage parasite samples (trophozoites and schizonts). This is due in part to the larger biomass of these stages but also to the existence of an effective enrichment method (7); erythrocytes infected with late stage parasites can be separated from uninfected erythrocytes using the paramagnetic properties of hemozoin, a byproduct of parasite maturation. The ability to enrich for late stages, thus limiting noise from excess uninfected erythrocytes, has fueled the recent explosion of omics-based studies of late stage *P. falciparum* biology and antimalarial drug action (8–16).

**Figure 1.**
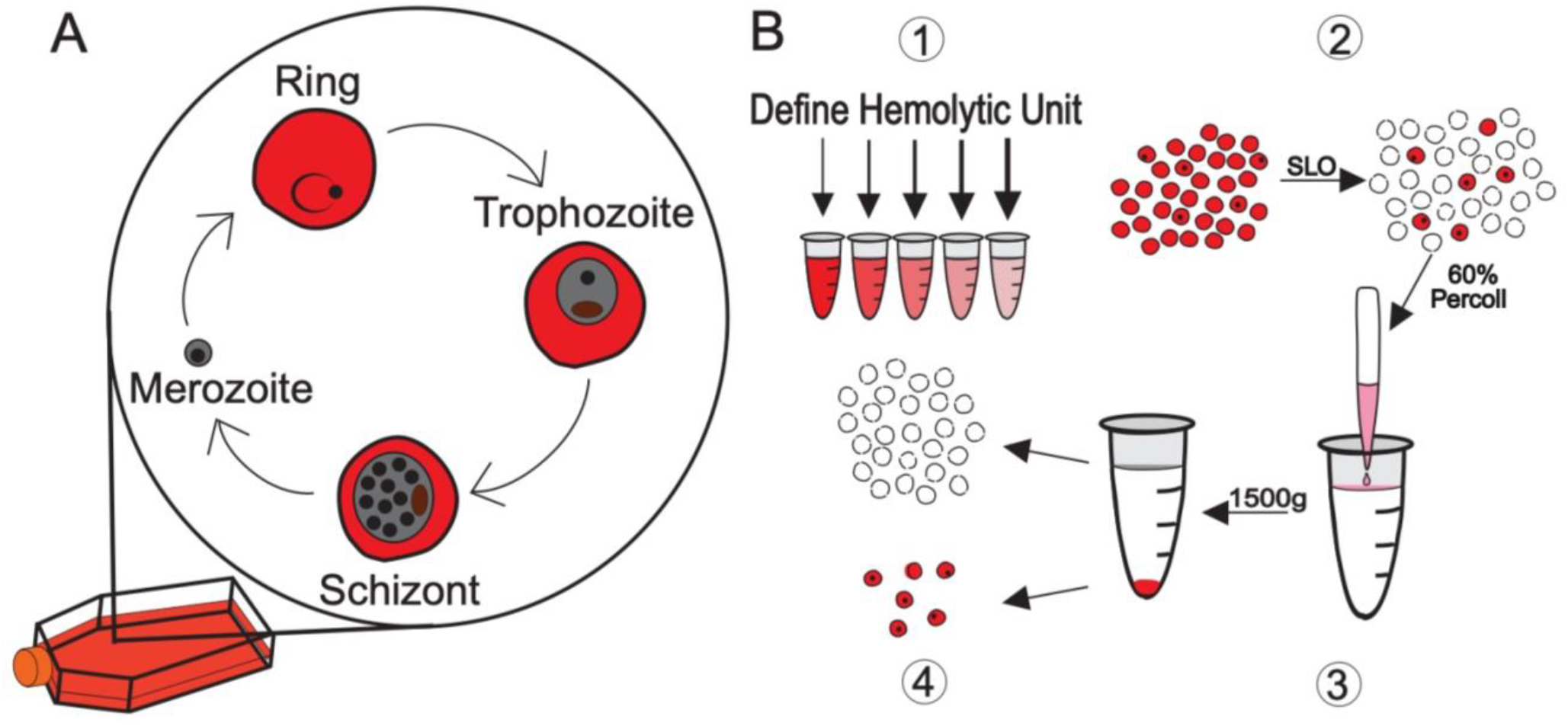
SLOPE enrichment overview. (A) The asexual replication cycle of *Plasmodium* occurs inside erythrocytes. Both *P. falciparum* and *P. knowlesi*, which take 48 and 24 hours, respectively, to complete the replication cycle, can be propagated *in vitro* indefinitely. (B1) Hemolytic activity of Streptolysin-O (SLO) was assessed on uninfected erythrocytes to define a unit (the amount of SLO necessary for 50% lysis of 50μl of uninfected erythrocytes at 2% hematocrit in PBS for 30 min at 37°C). (B2) Ring stage synchronized cultures were treated with a defined quantity of SLO units to preferentially lyse uninfected erythrocytes. (B3) SLO treated samples were layered over a 60% Percoll gradient and centrifuged to separate lysed ghosts from intact cells. (B4) The upper layer of Percoll containing lysed ghosts was discarded while the lower, intact, infected erythrocyte enriched fraction was collected. Uninfected erythrocytes, red circles; Infected erythrocytes, red circles with black dots; lysed membranous ghosts, white circle with dashed outline.

The lack of an effective enrichment method dramatically limits our ability to study the ring stage parasites, as well as ring-derived forms. For example, recent proteomics and metabolomics studies of this early erythrocytic *P. falciparum* show the heavy influence of host metabolites in non-enriched preparations, which contributes to variability between samples and obscures parasite phenotypes (16–18). This limitation is particularly acute when dealing with material directly from malaria patients as only ring forms of *P. falciparum* are collected during blood draws (19), and there is a high ratio of uninfected host cells to parasite-infected cells (typically 100:<4) (20). Ample access to clinically-relevant parasites is important for the study of antimalarial resistance especially in the context of artemisinin, which impacts rings differently than later stages through the induction of quiescence (21–23). This impactful biological discovery highlights the need to improve upstream purification steps for the study of ring stage biology.

In this study, we present a method for the enrichment of viable ring stage *Plasmodium-*infected erythrocytes that is simple to employ, rapid, and non-toxic to the parasite. This method can be scaled to the needs of individual experiments without compromising these attributes. We show that the effective removal of uninfected erythrocytes is unaffected by standard culture media formulations and is conserved across multiple *Plasmodium* species and parasite sources, further highlighting utility for a range of experimental needs. To our knowledge, this is the first enrichment method that is effective on ring stage parasites to increase parasitemia (the percentage of erythrocytes infected with a parasite) and reduce host erythrocyte contamination. The “SLOPE” enrichment method offers a tool to increase research quality in virtually all areas of *Plasmodium* asexual parasite research.

## Results

### A two-step SLOPE protocol effectively enriches ring-infected erythrocytes

We developed the Streptolysin-O (<u>SLO</u>)–<u>Pe</u>rcoll-based protocol (termed SLOPE) for enrichment of ring stage *P. falciparum* infected erythrocytes (Fig. 1B, see *Materials and Methods* and **Text S1** for protocol details). SLO is a pore forming toxin that preferentially lyses uninfected erythrocytes, leaving the large majority of infected erythrocytes intact (24). Using the protocol outlined by Jackson, *et al*. with slight modifications, we were able to achieve levels of lysis discrimination for erythrocyte populations that were comparable to this original report (93.4% and 9.9% lysis of uninfected and infected erythrocytes, respectively, Fig. 2A, yellow highlight). In addition to reproducing lysis levels, we quantified uninfected and infected erythrocyte lysis across a gradient of SLO concentrations. More complete lysis of uninfected cells (>99%) was obtained at the cost of greater infected erythrocyte lysis by increasing SLO quantity. For example, we found that 47U of SLO leads to >99% lysis of uninfected erythrocytes and 40% lysis of infected erythrocytes (Fig. 2A, grey highlight). Additionally, we show that SLO favors uninfected erythrocyte lysis irrespective of parasite line or culture media (Fig 2A; Fig S1). SLO showed comparable lysis discrimination between uninfected and infected erythrocytes in parasites grown in two common media formulations, RPMI 1640 supplemented with AlbuMAX II or RPMI 1640 supplemented with 20% human serum (Fig. 2A, maximum difference between uninfected and infected erythrocyte lysis: AlbuMAX supplementation = 83.5%; serum supplementation = 81.3%). Furthermore, SLO lysis showed considerable discrimination between uninfected and infected erythrocytes for both *P. falciparum* and the zoonotic species, *Plasmodium knowlesi* (Fig. 2A).

**Figure 2.**
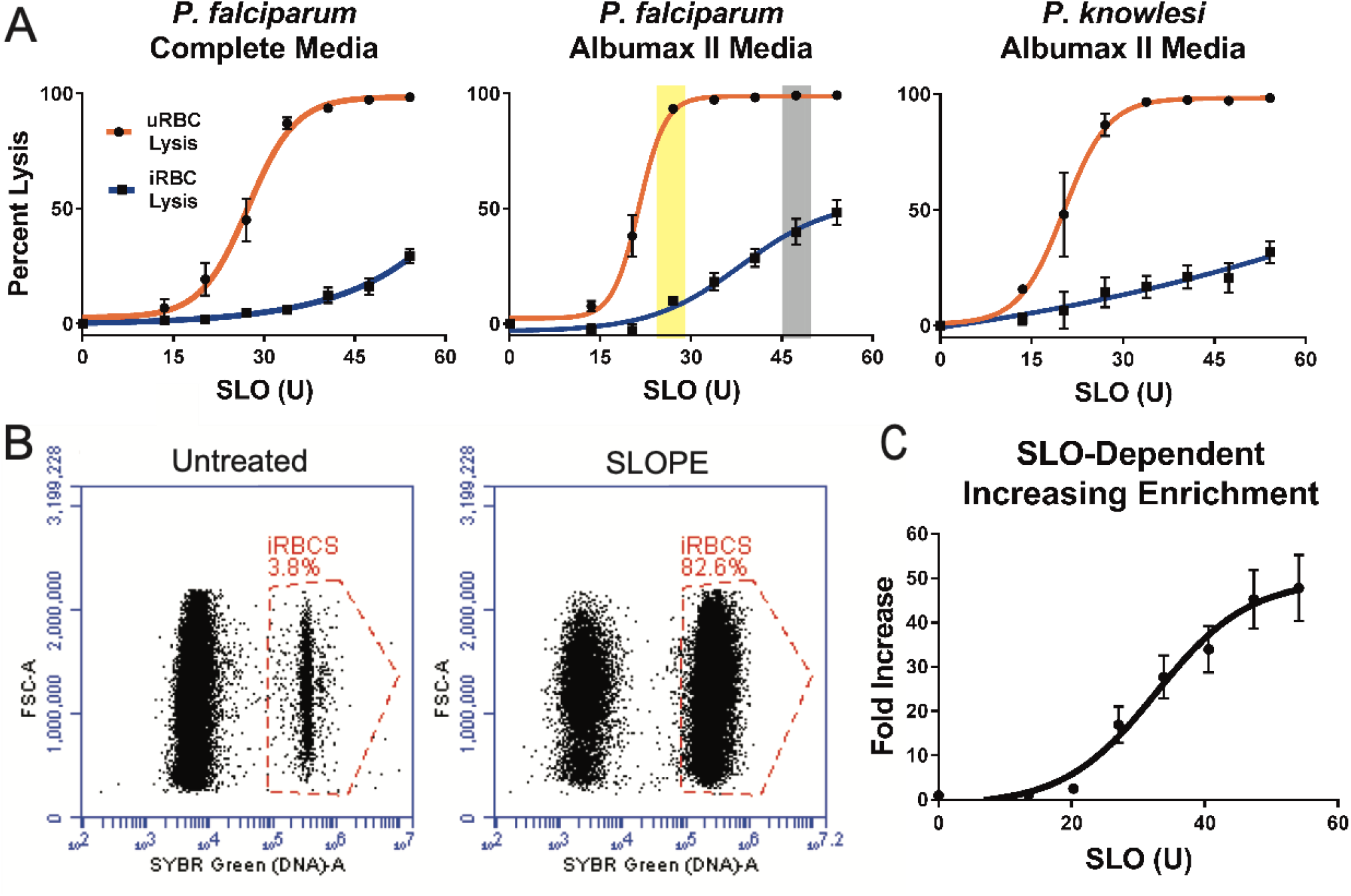
SLOPE enriches ring stage *Plasmodium* parasites irrespective of species or media formulation. (A) SLO lysis of uninfected erythrocytes (uRBCs) and infected erythrocytes (iRBCs) from (left) ring-stage synchronized *P. falciparum* grown in RPMI 1640 supplemented with 20% human serum (*N* = 6; 3 replicates each of lines Hb3 and K1), (middle) RPMI 1640 supplemented with Albumax II (*N* = 9; 3 replicates each of lines Hb3, K1, and MRA 1240), and asynchronous *P. knowlesi* grown in RPMI 1640 supplemented with Albumax II (*N*=3). (B) SYBR-Green based flow cytometry measurements before and after SLOPE enrichment. Flow plots show single cells within the erythrocyte size range. The infected erythrocyte fraction (“iRBCs”) is denoted within the dashed red gate. (C) Parasitemia fold increase upon treatment with increasing SLO units relative to untreated controls. Represented samples were grown in RPMI 1640 supplemented with Albumax II (*N* = 9; 3 replicates each of lines Hb3, K1, and MRA 1240). All error bars represent S.E.M.

After treatment with SLO, parasite samples contained a mixture of primarily lysed erythrocyte membranes (from uninfected erythrocyte lysis), termed ghosts, and intact erythrocytes (enriched in infected erythrocytes) (Fig. 1B2). For the second part of the enrichment protocol, we exploited the difference in density of intact erythrocytes and ghost membranes to separate the two populations (Fig. 1B3). We found that intact erythrocytes travel through a 60% Percoll gradient during centrifugation while erythrocyte ghosts remained above the Percoll layer (Fig. 1B4, Fig. 3). Collection of the intact erythrocyte population leads to a sample with up to a greater than 20-fold increase in the parasite to erythrocyte membrane ratio. In one representative trial, we demonstrate enrichment of ring stage samples to a final parasitemia over 80% (22-fold increase over starting parasitemia, Fig. 2B). Higher levels of enrichment were attainable with higher amounts of SLO (up to 48-fold with 55 units of SLO, Fig. 2C) but this is at the cost of parasite material (up to ~40% of infected erythrocytes, Fig. 2A).

**Figure 3.**
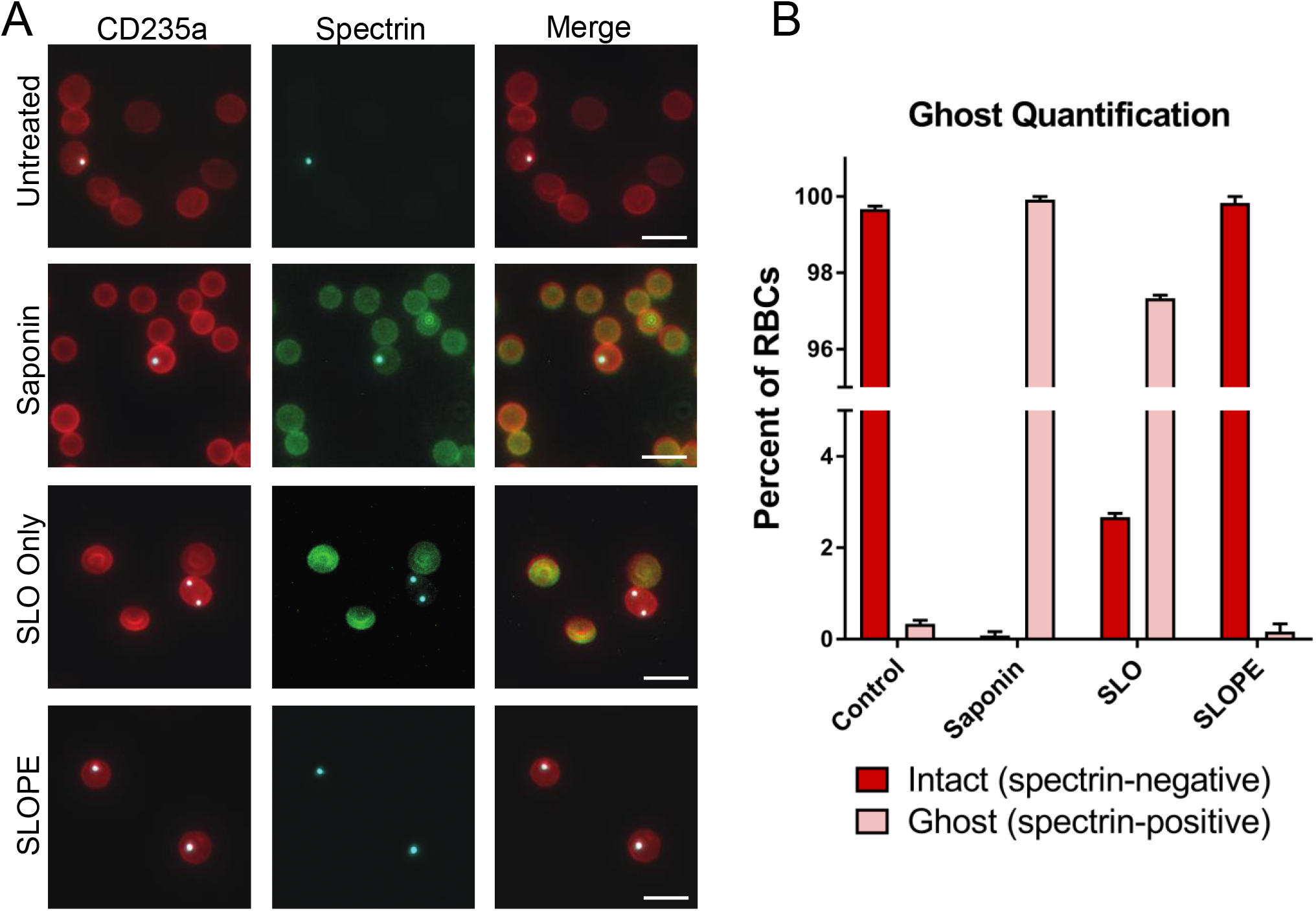
Validation of ghost separation from intact erythrocytes by Percoll step. (A) Intact erythrocytes are shown as CD235a (red) positive and spectrin (green) negative. Lysed erythrocytes, termed ghosts, are shown as CD235a and spectrin double positive (yellow in merge). All images show ring-stage *P. falciparum* line MRA 1240 parasites stained with SYBR Green (cyan). 40X Magnification; bar represents 10µm. Saponin samples were treated with 0.15% saponin for 5 minutes. SLO samples were treated with 40U of SLO for 6 minutes but were not centrifuged through a Percoll gradient. SLOPE samples were also treated with 40U SLO but were subjected to Percoll gradient centrifugation (B) Proportions of lysed ghosts and intact erythrocytes quantified from fluorescence microscopy imaging across different treatments using SYBR Green dye and CD235a and spectrin antibodies (*N* = 3; 400 erythrocytes per condition per preparation, error bars represent S.E.M.).

We performed SLOPE on mixed stage populations to determine whether our protocol biases enrichment of certain asexual stages. We found that SLOPE enrichment of lightly synchronized cultures (i.e. contains rings and some residual trophozoites) did not result in any alteration of the ring to trophozoite proportion (Fig. S2A). SLOPE enrichment was also unbiased when performed on asynchronous cultures (includes rings, trophozoites, and schizonts). However, the protocol did result in a non-significant drop in schizont percentage (Fig. S2B). Centrifugation and washes likely lysed fragile schizont-infected erythrocytes resulting in partial loss of the schizont population.

Next, we employed immunofluorescence detection of erythrocyte components to confirm that Percoll gradient density centrifugation removed ghost erythrocyte membranes from SLO treated samples (Fig. 3). Specifically, we visualized all erythrocyte membranes through detection of an external erythrocyte surface protein (CD235a); we identified ghosts through detection of an intra-erythrocyte protein (spectrin). Untreated (control) samples contain high erythrocyte membrane to parasite ratios (~100:1 due to ~1% parasitemia) as expected, and almost all membranes (>99%) were in the form of intact cells (spectrin negative, Fig. 3A and B). All erythrocyte membranes in samples treated with a nondiscriminatory lytic agent (saponin) were lysed ghosts (spectrin positive); however, the same high erythrocyte membrane to parasite ratio (100:1) that was observed in untreated samples persisted. In samples treated only with SLO (no Percoll gradient centrifugation), a subset of cells enriched for infected erythrocytes remained intact (1 of the 4 erythrocytes in Fig. 3A). However, the considerable lysed erythrocyte population remained, leaving the erythrocyte membrane to parasite ratio at 100:1. Complete SLOPE enrichment demonstrated the effectiveness of Percoll centrifugation for separating ghosts and intact cells by removing this lysed erythrocyte population, leaving a >99% intact erythrocyte population that is enriched for parasites (25-fold decrease in the erythrocyte membrane to parasite ratio; starting of 100:1, final of 4:1). Image quantification revealed that <0.2% of erythrocytes in SLOPE enriched samples were ghosts (Fig. 3B).

### Parasites retain full viability after SLOPE enrichment

We sought to test the effect of SLOPE enrichment on parasite metabolism and viability. First, parasites were stained with a mitochondrial membrane potential dependent dye to determine the fraction of metabolically active parasites. The percentage of parasites with an active mitochondrial membrane potential was not significantly different between SLOPE enriched samples and untreated controls (*untreated*=97%, *SLOPE*=94%, *p*=0.46, Fig. 4A). To assess parasite viability on a longer-term scale, we monitored parasite growth following SLOPE enrichment. This was accomplished by diluting SLOPE enriched parasites with fresh erythrocytes to reduce parasitemia to appropriate levels for culture (<5%). When compared to that of parasitemia-matched non-enriched, untreated flasks, the growth rate of parasites from SLOPE enriched samples showed no growth defects over the time period measured (6 days) (Fig. 4B). Ring stage parasites retained normal morphology following SLOPE enrichment, further providing evidence that SLOPE enrichment did not damage parasites (Fig. 4C).

**Figure 4.**
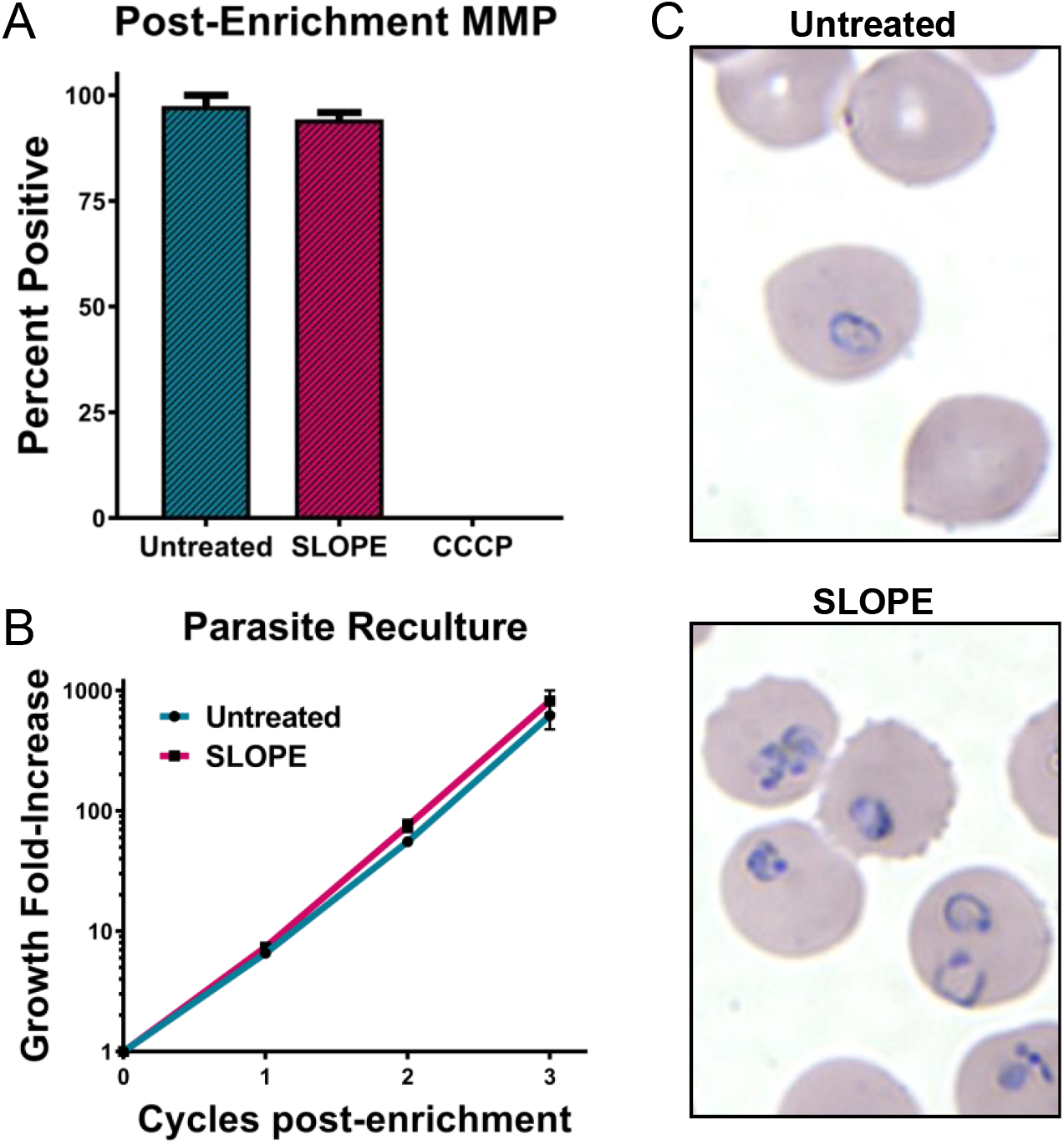
SLOPE enriched parasites remain viable. (A) Mitoprobe DiIC1 (5) mitochondrial membrane potential (MMP) measurements obtained by flow cytometry in untreated and SLOPE enriched ring-stage *P. falciparum* line MRA 1240 parasites (*N* = 3, error bars represent S.E.M.). (B) Six day of *P. falciparum* line MRA 1240 parasite growth from untreated controls or SLOPE enriched samples diluted with uninfected erythrocytes (*N* = 4, error bars represent S.E.M.). (C) Ring-stage *P. falciparum* line Dd2 infected erythrocytes visualized by Giemsa-stain at 100X magnification.

### SLOPE-enriched samples exhibit increased parasite metabolites

We hypothesized that a reduction of excess erythrocyte ghosts following SLOPE will lead to increased parasite signal, especially in ring stage samples. In order to investigate this impact, we compared synchronous ring stage *P. falciparum* parasites that were either untreated or SLOPE enriched by 15-fold using a plate-based targeted metabolomics platform that detects and quantifies up to 180 defined small molecules (*N*=4 per condition, 3×10_8_ erythrocytes per replicate) (Fig. 5A). One hundred and sixteen metabolites were detected in at least 50% of samples, representing 11 acylcarnitines, 17 amino acids, 5 polyamines, 69 glycerophospholipids, and 14 sphingolipids. Principal component analysis revealed distinct separation of untreated and SLOPE samples indicating a clear contribution of the increased concentration of parasite metabolites (Fig. 5B). In general, lipids contribute heavily to the separation of untreated and SLOPE groups with phosphatidylcholines having a large impact on the metabolic differentiation of the two groups (Table S1), which is in line with the observation that this lipid class is dramatically increased in infected erythrocytes (25).

**Figure 5.**
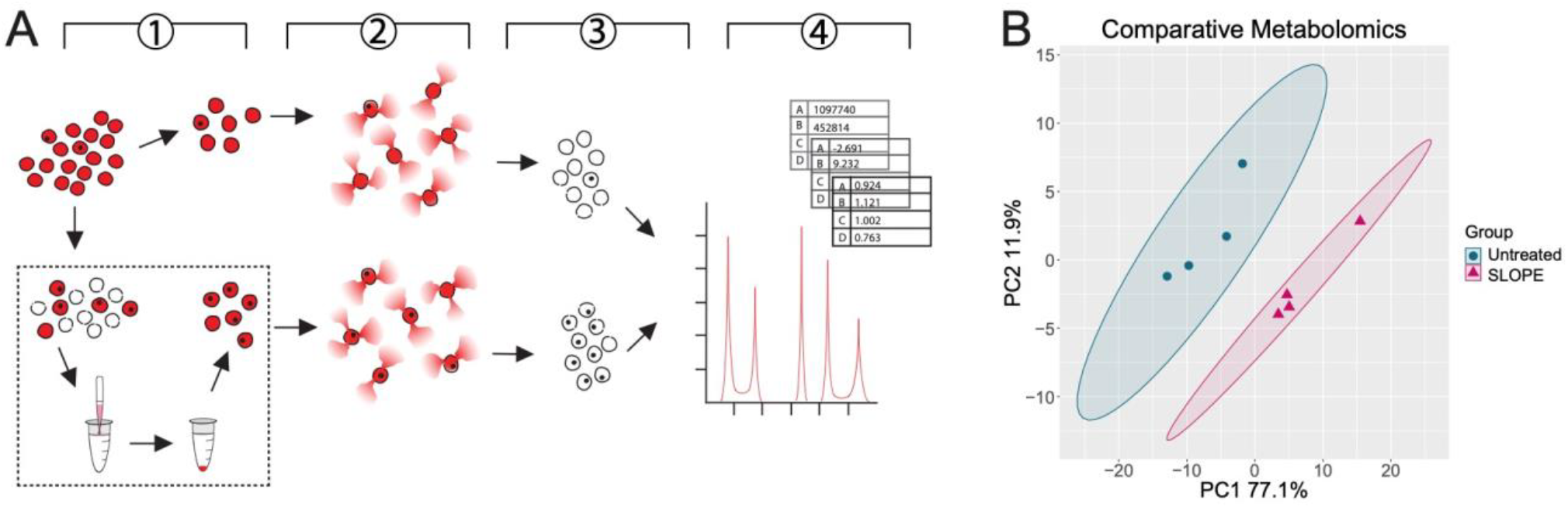
SLOPE enrichment increases detection of ring stage parasite metabolome. (A) Pipeline outlining metabolomics sample preparation and analysis. 1) Ring-stage synchronized *P. falciparum* Dd2 cultures were split into two fractions: one portion was taken for the untreated group and one portion was SLOPE enriched. 2) Equal numbers of erythrocytes from untreated and SLOPE groups were saponin lysed and washed to remove cytosolic erythrocyte metabolites. 3) Metabolism of the resulting pellet containing erythrocyte ghosts and parasites was quenched and metabolites were extracted. 4) Metabolites were identified and quantified from extracts using the AbsoluteIDQ p180 kit. These data were log transformed, centered, and scaled prior to statistical analysis. (B) Principal component analysis was performed on all metabolites detected in at least 50% of samples. Significance between untreated and SLOPE groups was determined by perMANOVA: *p*=0.037. Ellipses show 95% confidence intervals.

Thirty-two metabolites were significantly different between untreated and SLOPE groups after multiple testing correction (Table 1). While some conservation of metabolites was expected due to the persistence of erythrocyte membranes, metabolites that significantly differed between groups reflected expected biological differences associated with increased parasite signal. SLOPE enriched samples exhibited a substantial increase in 12 glycerophospholipids and 3 sphingolipids (3.13 and 5.64 average fold change, respectively, Table 1), which was likely due to the contribution of parasite membranes and other lipid-containing structures in parasitized erythrocytes (26). Twelve amino acids were higher in SLOPE samples (14.8 average fold change, Table 1), indicative of the breakdown of hemoglobin by the parasite into free amino acids (27). Additionally, metabolites of the polyamine synthesis pathway, which is active in the parasite (28, 29), were dramatically increased (10.3 average fold change, Table 1).

**Table 1.** SLOPE enrichment leads to significant differences in detection of the ring stage metabolome.

\*Denotes metabolites in the polyamine synthesis pathway.
| Metabolites | Adjusted p-value | SLOPE/Control |
| --- | --- | --- |
| *Polyamines* | | $\bar{x} = 10.3$ |
| Putrescine* | 0.000848 | 13.9 |
| Spermidine* | 0.000848 | 11.5 |
| Spermine* | 0.00796 | 5.53 |
| *Amino Acids* | | $\bar{x} = 14.8$ |
| Alanine | 0.0296 | 13.2 |
| Arginine* | 0.0204 | 6.94 |
| Asparagine | 0.0125 | 12.3 |
| Aspartic Acid | 0.0380 | 20.3 |
| Glutamic Acid | 0.0204 | 25.0 |
| Lysine | 0.0380 | 3.14 |
| Ornithine* | 0.0287 | 2.73 |
| Phenylalanine | 0.0204 | 8.11 |
| Serine | 0.0485 | 8.92 |
| Threonine | 0.0204 | 17.3 |
| Tyrosine | 0.0204 | 20.8 |
| Valine | 0.0170 | 38.7 |
| *Glycerophospholipids* | | $\bar{x} = 3.13$ |
| lysoPC.a.C16:0 | 0.0485 | 1.70 |
| lysoPC.a.C17:0 | 0.0313 | 1.40 |
| lysoPC.a.C18:1 | 0.0249 | 2.67 |
| lysoPC.a.C18:2 | 0.0204 | 3.74 |
| PC.aa.C32:0 | 0.0290 | 5.91 |
| PC.aa.C36:1 | 0.0380 | 5.29 |
| PC.aa.C38:5 | 0.0485 | 3.10 |
| PC.aa.C40:4 | 0.0289 | 3.57 |
| PC.ae.C38:0 | 0.0380 | 2.36 |
| PC.ae.C38:3 | 0.0287 | 2.61 |
| PC.ae.C40:1 | 0.0447 | 2.79 |
| PC.ae.C40:4 | 0.0204 | 2.40 |
| *Sphingolipids* | | $\bar{x} = 5.64$ |
| SM.C24:0 | 0.0290 | 7.44 |
| SM.OH.C24:1 | 0.0380 | 4.58 |
| SM.C24:1 | 0.0445 | 4.91 |
| *Acylcarnitines* | | $\bar{x} = 2.19$ |
| C2 | 0.0380 | 3.78 |
| C16 | 0.0447 | 0.60 |

### SLOPE enriches clinical samples in a cholesterol dependent manner

In order to determine the utility of SLOPE for the enrichment of clinical *P. falciparum* parasites, which predominantly circulate as ring stages, we tested SLO lytic discrimination between infected and uninfected erythrocyte populations directly from human patients. Contrary to results with *in vitro* propagated parasites (Fig. 2A), patient erythrocytes demonstrated reduced SLO lytic discrimination (Fig. 6A, left panel). However, short-term (6h) incubation of clinical samples in complete media (RPMI with 20% human serum) greatly increased SLO lytic discrimination, thereby decreasing parasite loss leading to a higher possible final parasitemia (Fig. 6A, right panel). Restoring SLO discrimination occurred with as little as 4h of *ex vivo* incubation but the benefit peaked at ~6h (Fig. S3).

**Figure 6.**
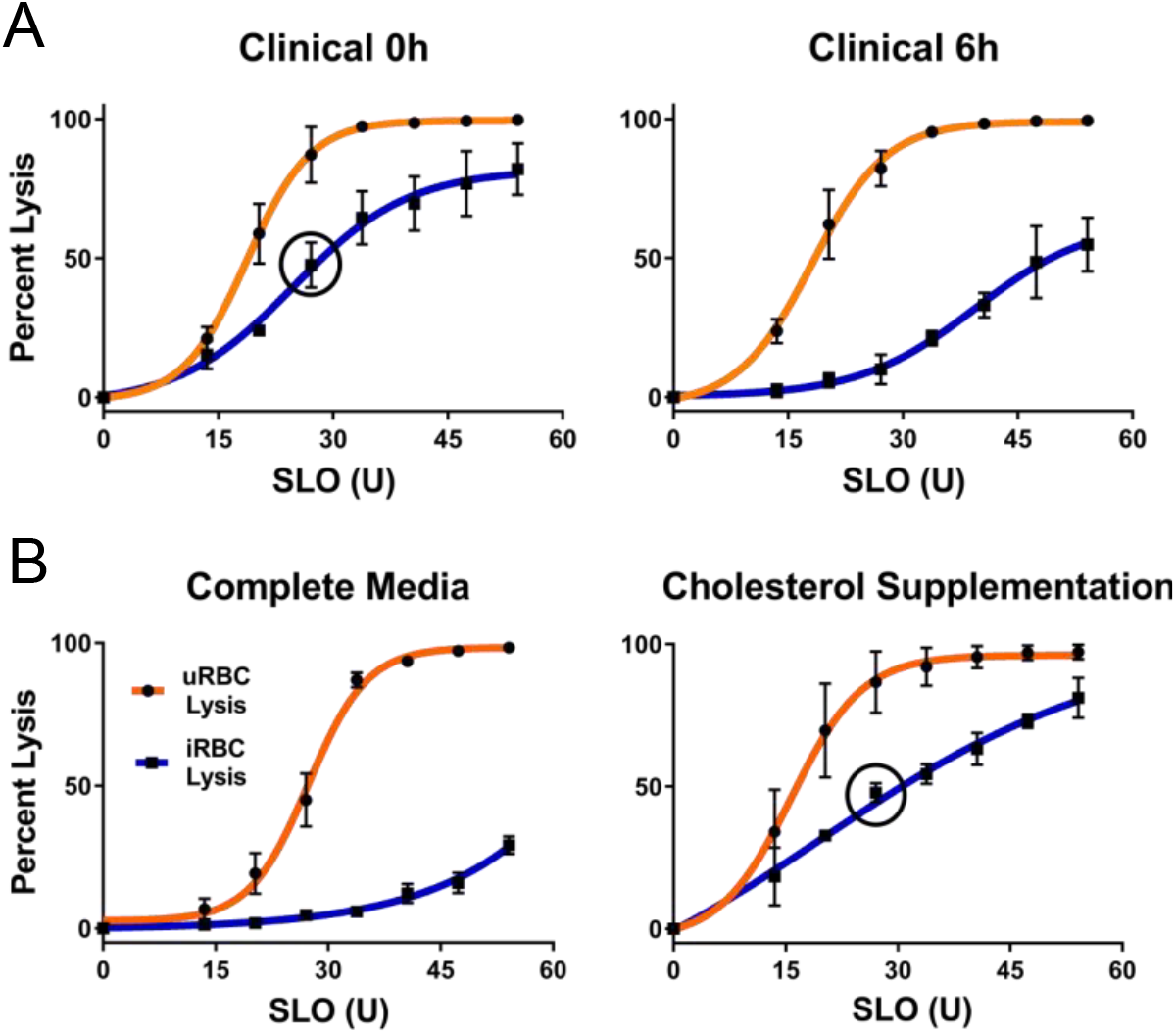
SLOPE is effective on clinical samples in a cholesterol dependent manner. (A) SLO lysis of erythrocytes from *P. falciparum* infected patients either directly isolated from the patient (0h) or after 6h of incubation in complete media (RPMI supplemented with 20% serum). *N*=3 patients. Error bars represent S.E.M. (B) SLO lysis of laboratory *P. falciparum* either grown for 48h in complete media or complete media supplemented with 4mM cholesterol saturated mβCD. *N*=6 (left; 3 replicates each of lines Hb3 and K1); *N*=3 (right; line Hb3). Error bars represent S.E.M. Black circles in selected graphs show iRBC lysis at 27U SLO.

We hypothesized that the reduced SLO lytic discrimination observed in samples upon immediate acquisition from patients was due to the increased abundance of cholesterol *in vivo* compared to *in vitro* media formulations. As expected, the *in vivo* environment contained cholesterol at concentrations up to ~250 times higher than those found *in vitro* (Table S2). To test the exchange of cholesterol between serum and erythrocyte membranes, we compared the SLO lysis of laboratory *P. falciparum* grown in complete media to parasites grown in complete media supplemented with cholesterol (Fig. 6B). This artificial addition of cholesterol led to the predicted increase in infected erythrocyte SLO lysis, closely mimicking the reduced SLO lytic discrimination observed upon immediate acquisition of clinical samples from patients (47.5% lysis and 47.8% lysis in clinical *ex vivo* and cholesterol supplemented *in vitro* samples, respectively, when treated with ~30U SLO; Fig. 6, black circles).

### SLOPE enriches in vitro-derived quiescent parasites

A low frequency of ring stage *P. falciparum* parasites has been reported to enter a quiescent state upon artemisinin treatment. In order to explore the effectiveness of our enrichment method on non-traditional erythrocytic forms of the parasite, we treated parasites with dihydroartemisinin (DHA), removed actively growing parasites daily, and then performed SLOPE to enrich the remaining metabolically quiescent parasites. When quiescent parasites were quantified using flow cytometry (SYBR-Green positive/MitoProbe positive), we detected an enrichment in quiescent parasites by >100-fold (pre-enrichment: undetectable but estimated based on previous studies to be ~0.005 quiescent parasites/100 erythrocytes (30); post-enrichment: 1 quiescent parasite/100 erythrocytes). Quiescent parasites were also readily visually identifiable in SLOPE enriched samples under microscopy (Fig. 7A, SYBR-Green/MitoTracker Red positive). When compared to untreated rings, quiescent parasites showed reduced MMP staining area, indicative of the condensed cytoplasm and reduced metabolism seen in quiescence (31, 32). While quiescent parasites were enriched by the removal of uninfected erythrocyte material, erythrocytes containing dead parasites were also enriched by this process (Fig. 7A and B, SYBR-Green positive, Mitotracker/Mitoprobe negative parasites). However, live, quiescent parasites appeared to be enriched at considerably higher rates compared to their dead counterparts. Specifically, the dead parasite population experienced less than 10-fold enrichment (1.2% immediately pre-SLOPE versus 9.1% post-SLOPE), whereas mentioned above, quiescent-infected erythrocytes were enriched by >100-fold. Overall, ~one-tenth of total enriched parasites (SYBR-Green positive events) were quiescent (Mitoprobe positive events) (Fig. 7B), despite estimates that <<1% of all parasites enter quiescence upon DHA treatment (30).

**Figure 7.**
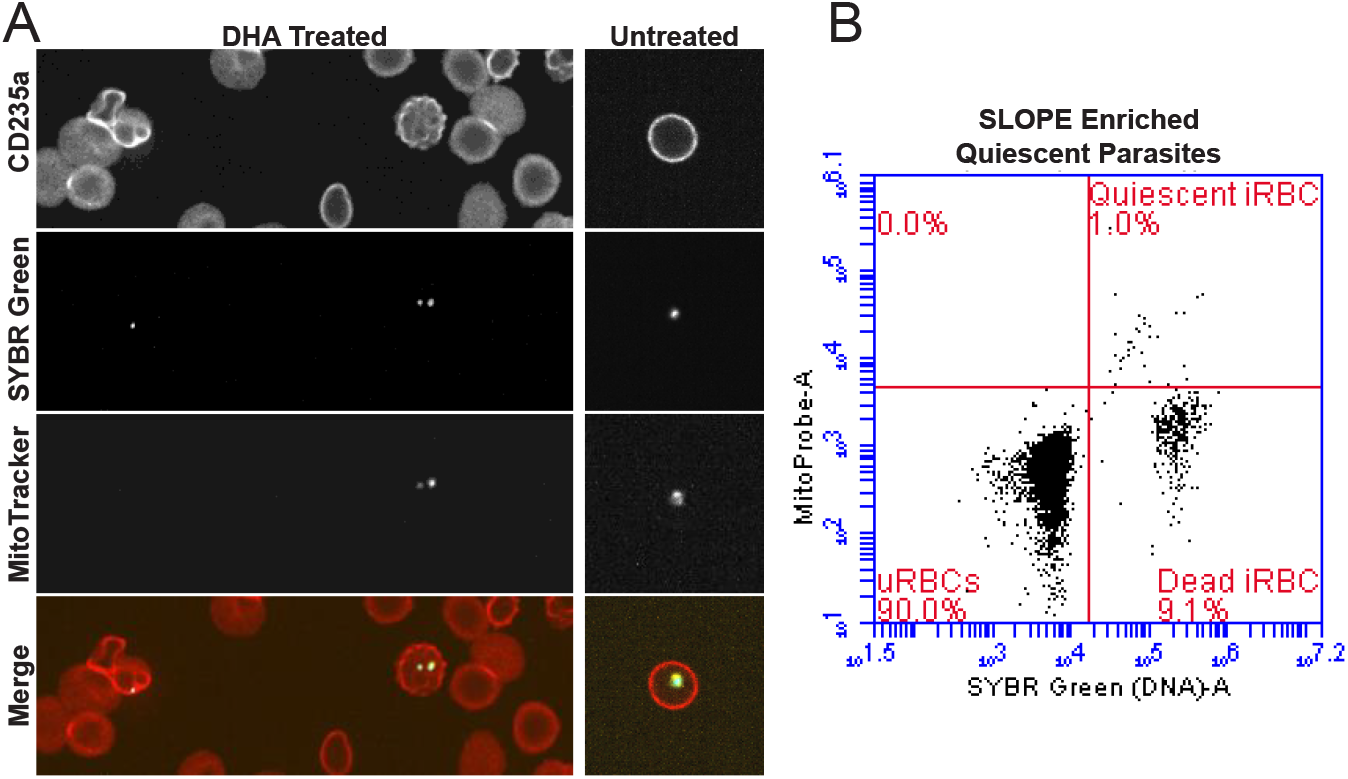
SLOPE is effective on DHA-induced quiescent parasites. (A) Erythrocytes are shown by CD235a staining and *P. falciparum* MRA 1238 parasites are shown by SYBR Green. Within the DHA-treated image, a dead parasite (left) failed to accumulate MitoTracker Deep Red, while two quiescent parasites accumulated MitoTracker Deep Red. (B) Flow cytometry plot measuring quiescent parasites as SYBR Green and Mitoprobe double positive events. Dead parasites are SYBR Green positive, but Mitoprobe negative; uninfected erythrocytes (uRBCs) are SYBR Green and Mitoprobe double negative. iRBC = infected erythrocyte.

## Discussion

For the first time, we can achieve considerable ring stage parasite enrichment that is effective across a variety of conditions and *Plasmodium* species (Fig. 1; Fig. 2), as well as on understudied ring-related populations (clinical isolates, Fig. 6; quiescent forms, Fig. 7). Our two-part SLOPE enrichment relies on the cholesterol-dependent lysis of uninfected erythrocytes, followed by the exploitation of density differences between a parasite-rich intact population and a ghost population of primarily uninfected erythrocyte membranes (Fig. 3; Fig. 6). Importantly, resulting enriched infected erythrocytes maintain active metabolism and growth (Fig. 4; Fig. 5; Table 1).

The SLOPE method expands on a discovery made over a decade ago (24) and integrates our metabolic knowledge about the parasite. *Plasmodium* species lack the ability to synthesize cholesterol *de novo* and therefore, the parasite scavenges this lipid from host erythrocyte membranes (33). This scavenging likely leads to the lower levels of cholesterol observed in infected erythrocyte plasma membranes (26). Upon discovery of the preferential action of SLO on uninfected erythrocytes, reduced levels of cholesterol in the membranes was proposed as the mechanism of discrimination (24). Our study on parasites from a cholesterol rich environment and the impact of cholesterol addition on SLOPE effectiveness add validity to this hypothesis (Fig. 6B). We showed that when parasites were isolated from the human patient (100% serum), SLO discrimination was nominal. Since cholesterol is readily exchanged between the serum environment and erythrocyte membranes (34), we predict that this result is due to the replenishment of parasite-scavenged cholesterol in the erythrocyte membrane (Fig. 8). While enrichment of parasites directly from the bloodstream of a patient is possible, this is at the expense of parasite number (Fig. 6A, Fig. S3). Fortunately, parasite loss can be minimized with a short-term incubation in the laboratory to reduce erythrocyte membrane cholesterol levels; the limited duration (≤6h) and use of 20% human serum during this step will likely allow parasites to maintain *in vivo* qualities (e.g. transcription and metabolic program). Further studies will be required to determine if this is the case.

**Figure 8.**
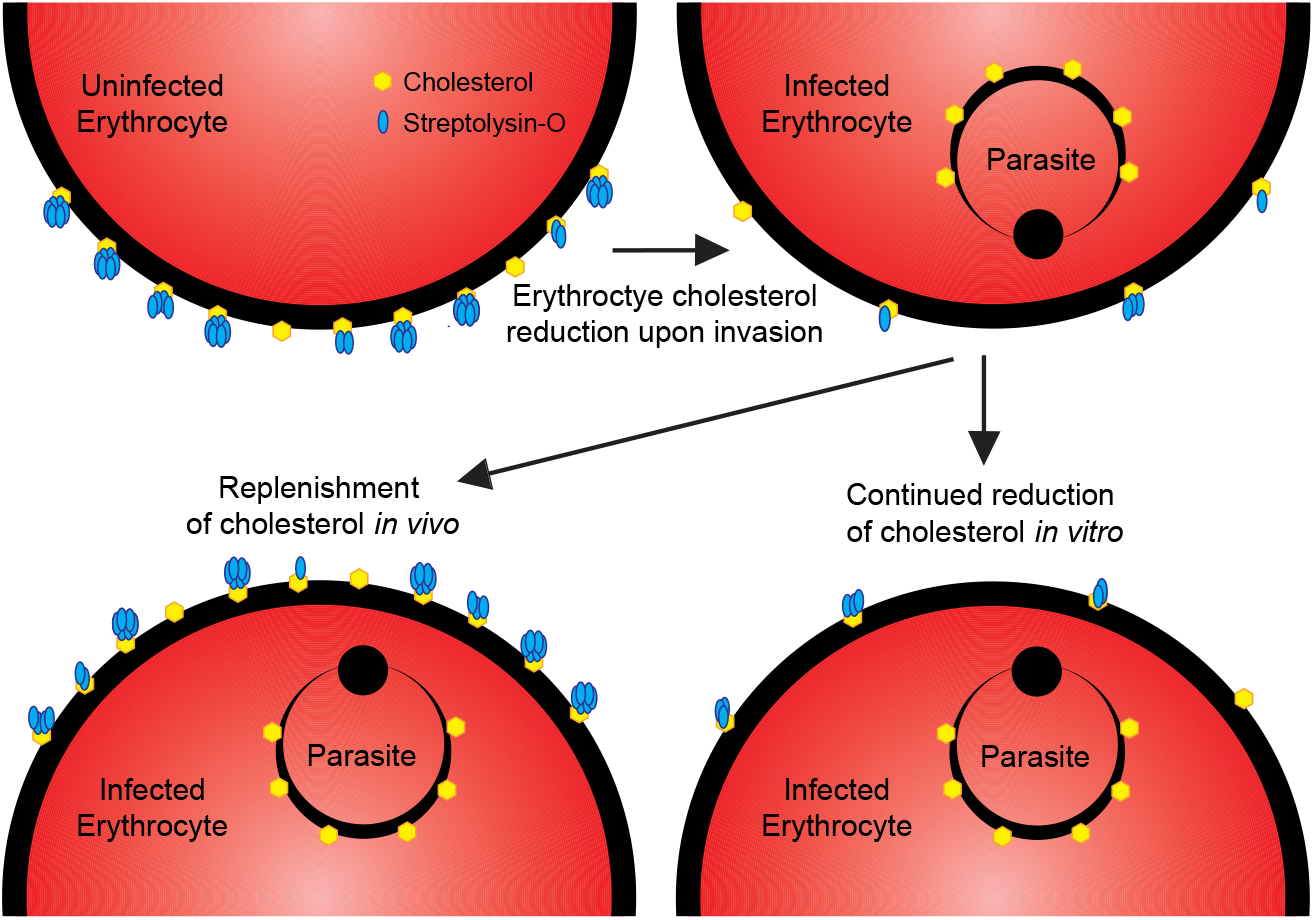
Proposed mechanism for decreased susceptibility of infected erythrocytes to SLO lysis. Upon invasion of an erythrocyte, the parasite salvages host membrane cholesterol leading to lower cholesterol levels on the infected erythrocyte surface (top of diagram). During *in vitro* incubation in conditions with sub-physiological cholesterol levels, cholesterol remains low via continued parasite scavenging. Upon exposure physiological levels of cholesterol, such as *in vivo*, the exchange of cholesterol between plasma and erythrocytes restores erythrocyte cholesterol to near pre-invasion levels.

SLOPE shows utility for the enrichment of non-traditional populations of *Plasmodium* parasites that were previously understudied due to limited accessibility. In previous studies, clinical parasites were often fully adapted to *in vitro* culture to generate enough material for characterization (35, 36). This is not ideal considering parasite populations are altered by the selective pressure and environmental changes upon transition to *in vitro* culture (37–39). Further, a sizeable portion of clinical isolates fail entirely to adapt to *in vitro* conditions (40). With the direct enrichment of parasites from patients, many studies can now be performed with minimal perturbations (see above) or without culture adaptation entirely.

Another form of *P. falciparum* that remains understudied is the artemisinin-induced quiescent state that is somewhat analogous to persister cells seen in bacteria (41). Populations of both quiescent parasites and bacterial persisters demonstrate arrested growth leading to decreased susceptibility in the face of stressful conditions, such as drug treatment. Quiescent and persister metabolic states share similarities as well, including downregulation of DNA replication, tRNA synthesis, and oxidative metabolism but maintenance of fatty acid synthesis (42). Following stressor removal, both populations have the ability to resume growth leading to a recrudescent infection. Parasite recrudescence following quiescence presents a challenge to the efficacy of antimalarial therapies. Yet, the rarity of quiescent parasites following drug treatment (<<1% of all parasites, (30)) limits the study of this phenomenon. Excitingly, SLOPE enrichment increases the parasitemia of quiescent parasites to readily detectable levels (~1%) through both the reduction of uninfected erythrocytes and the partial reduction of erythrocytes containing dead parasites (Fig. 7). At detectable levels, it is feasible to further enrich quiescent parasite populations using other methods (i.e. fluorescence activated cell sorting). The increased enrichment preference for live, quiescent parasites over dead parasites is likely due to the loss of cholesterol metabolism in parasites upon death. While quiescent parasites down-regulate many aspects of metabolism (31), these data indicate that quiescent parasites continue the utilization of cholesterol.

The utility of SLOPE enrichment in increasing access to both *in vitro* rings and non-traditional parasite forms (quiescent parasites and clinical rings) will reduce many of the previous barriers to high-quality *Plasmodium* research. Lack of an effective enrichment method for ring stage *Plasmodium* has led to considerable host-contributed noise in samples, limiting the success of sensitive downstream analyses such as proteomics and metabolomics. Specifically, our previous work demonstrated that characteristics of the erythrocyte batch contributed to the resultant ring parasite metabolome as heavily as drug treatment (17). Additionally, erythrocyte noise stymied previous ring stage metabolomics and proteomics experiments as parasite-derived signals could not reliably be determined over the host erythrocyte contribution (16–18). Since parasitemia was low in these studies (~1-5 parasites per 100 erythrocytes), this was undoubtedly due to the high number of erythrocyte ghost membranes that remained in the preparations. SLOPE significantly reduces the number of ghosts in our preparations, thus drastically increasing parasitemia (up to >80% in some cases, Fig. 2B). We directly observed this increase in parasite specific metabolites over the host erythrocyte noise using targeted metabolomics (Fig. 5B, Table 1). Most notably, we detected an increase in metabolites of the polyamine synthesis pathway that was proportional to the parasite enrichment level (mean of ~10-fold and 15-fold, respectively). Uninfected erythrocytes lack the machinery necessary for polyamine synthesis and contain only low levels of putrescine, spermidine, and spermine (43). However, polyamine levels are much higher in proliferating cell, such as *Plasmodium*, which both scavenge and synthesize polyamines (28, 29). Given the association between parasite number and polyamine concentration, we propose to use this group of metabolites as a quality control marker for parasite-derived metabolites in future metabolomics investigations.

The SLOPE method has several advantages for the isolation of material to be used for sensitive downstream analyses. Firstly, our method can be performed rapidly (~30-40 min). This is equal to or faster than the amount of time needed to perform the field-accepted purification method for isolation of late stage parasites by magnetic column purification (~30min to 1hr). The rapid purification ensures the cellular state of samples is as close as possible to the true phenotype. Secondly, this method is highly scalable and can be used on culture amounts that range from a fraction of a milliliter to combined pools of many flasks (hundreds of milliliters). Finally, SLOPE enrichment is facile and only requires access to a few reagents and a standard centrifuge. While a fluorescence-based flow cytometer and an automated brightfield cell counter were used to perform the experiments in this paper, less costly devices, such as a microscope and hemocytometer, respectively, can be used. The addition of SLOPE to our laboratory toolset will be useful not only for proteomic and metabolomics studies (see above) but also genomic and transcriptomic analyses. Reduction of contaminating host DNA and RNA that remain associated with erythrocytes relative to parasite material will lead to increased sequencing coverage particularly in clinical samples, which contain a considerable human contribution (44).

One caveat of this method may be the cost of SLO lytic agent. Researchers seeking to regularly enrich large quantities of parasites may consider in-house isolation of SLO from *Streptococcus pyogenes* culture or recombinant SLO expression in *Escherichia coli* in place of purchasing commercially available SLO. An additional caveat of SLOPE enrichment is the loss of parasite material. However, due to the scalability of this method, starting material can be increased to compensate for this loss. When sample material is limited, researchers may wish to use lower quantities of SLO to minimize lysis of infected erythrocytes.

With SLOPE enrichment as a tool, researchers can more accurately study the ring stage of the *Plasmodium* erythrocytic cycle. Improvements in our understanding of parasite biology will lead to more effective treatments for this deadly disease. Beyond this use, SLOPE has the potential to be extended to other intracellular parasites and cell types. Many Apicomplexan parasites scavenge cholesterol from host cells. For example, asexual replication of *Toxoplasma gondii* within nucleated cells requires cholesterol scavenging with cholesterol being transferred from the host cell membrane to the parasitophorous vacuole immediately upon invasion (45). This presents the possibility that other cells types infected with cholesterol scavenging parasites will also display increased resistance to SLO lysis. Further, SLO causes cholesterol dependent membrane lysis in cell types other than erythrocytes (46, 47). Thus, the SLOPE method shows potential for the enrichment of cells infected with other intracellular, cholesterol-scavenging pathogens, including *Toxoplasma*, *Cryptosporidium*, *Theileria, Chlamydia*, and *Mycobacterium*.

## Materials and Methods

### Parasites and growth

*Plasmodium falciparum* lines (MRA-155 (Hb3), MRA-159 (K1), MRA 150 (Dd2), MRA 1240, and MRA 1238, use of respective lines are indicated in figure legends) were obtained from the Malaria Research and Reference Reagent Resource Center (MR4, BEI Resources); The human erythrocyte-adapted *Plasmodium knowlesi* line (yH1, (48)) was a gift from Manoj Duraisingh (Harvard University T.H. Chan School of Public Health). *Plasmodium* cultures were maintained in A+ human erythrocytes at 3% hematocrit in RPMI 1640 HEPES (Sigma Aldrich, St Louis, MO) supplemented with either 0.5% Albmumax II Lipid-Rich BSA (Sigma Aldrich, St Louis, MO) and 50 mg/L hypoxanthine (Thermo Fisher Scientific, Waltham, MA) or with 20% heat inactivated human serum (cRPMI) (Valley Biomedical, Winchester, VA). Cultures were grown at 37°C under 5% oxygen, 5% carbon dioxide, 90% nitrogen gas (49). Dilution of cultures with uninfected erythrocytes to maintain parasitemia at <5% was performed every other day as was changing of culture medium. Parasitemia was determined by flow cytometry using SYBR-Green staining (50). Cultures were confirmed negative for mycoplasma monthly using a LookOut Mycoplasma PCR detection kit (Sigma Aldrich, St Louis, MO).

Parasites from infected patients were obtained from adults admitted to the University of Virginia Health System with clinical malaria. All patients had a recent history of travel to a malaria-endemic African country, and *P. falciparum* infection was clinically determined by a positive rapid diagnostic test and peripheral blood smear analysis. Samples were obtained within 24h of phlebotomy and washed twice with RPMI 1640 HEPES to remove white blood cells. Erythrocytes were then either immediately used for experiments or kept under short-term *in vitro* culture conditions in cRPMI. All samples were handled in accordance with protocols approved by the University of Virginia Institutional Review Board for Health Sciences Research.

### SLO activation and storage

Streptolyisn-O (SLO, Sigma Aldrich, St Louis, MO) activation was performed as previously described (24). Following activation, SLO was stored in aliquots at −20°C until use. Hemolytic units (U) of each SLO batch were quantified from a stored aliquot; one unit equals the amount of SLO necessary for 50% lysis of 50μl uninfected erythrocytes at 2% hematocrit in PBS for 30 min at 37°C. Hemolytic activity was recurrently assessed (approximately monthly) in triplicate to control for SLO degradation over time and varying cholesterol levels contributed by different media batches.

### SLOPE enrichment

When required, *P. falciparum* cultures were synchronized to the ring stage prior to enrichment using one round of 5% D-sorbitol (Sigma Aldrich, St Louis, MO) (52). SLO lysis was performed as previously described but with modifications (24) (Fig. 1B; **Text S1**). Briefly, erythrocyte density was measured using a Cellometer Auto T4 (Nexcelom Biosciences, Lawrence, MA). Cell density was adjusted to 2×10_9_ erythrocytes/mL. The desired amount of SLO (between 0U and 55U) was added in a ratio of 2 parts SLO solution to 5 parts erythrocytes. Samples were mixed well by pipetting and incubated at room temperature for precisely 6 min. Five-10 volumes of 1X PBS or non-cholesterol containing media (ex. RPMI 1640 HEPES) were added and cells were centrifuged at 2,500xg for 3 min. After removal of the supernatant, cells were washed twice more with 1X PBS or non-cholesterol containing media. Following SLO lysis, cells suspended in 1X PBS were layered onto a 60% Percoll gradient (Sigma Aldrich, St Louis, MO) and centrifuged at 1,500xg for 5-10 min depending on the volume of the gradient. The top layer of Percoll was discarded while the lower cell pellet was collected and washed twice with 1X PBS or media.

### SLO lysis curves

*Plasmodium* were treated with a range of SLO units as described above. Flow cytometry was used to assess equal volumes of each sample for every experiment on a BD Acurri C6 flow-cytometer (BD Biosciences, San Jose, CA.). Total erythrocyte values were obtained by gating for intact erythrocytes based on forward and side scatter. Infected erythrocyte values were obtaining by gating for SYBR-Green positive intact erythrocytes. Uninfected erythrocyte values were obtained by subtracting infected erythrocyte values from total erythrocyte values. Percent lysis of uninfected and infected erythrocyte populations were determined by comparing the infected and uninfected erythrocyte values for each SLO-treated sample in an experiment to the untreated control.

### Ghost Quantification

CD235a is constitutively present on the erythrocyte outer surface and thus, both intact erythrocytes and lysed ghosts are CD235a+. Conversely, spectrin is located in the erythrocyte cytoskeleton meaning anti-spectrin cannot access the antibody target in intact erythrocytes. However, pores in lysed erythrocyte membranes (saponin or SLO treated) allow passage of antibodies into the erythrocyte cytosolic compartment making these cells spectrin+. For fluorescent imaging, unfixed samples (average parasitemia of 1%) were blocked with 2% BSA followed by staining in suspension with a 1:100 dilution of mouse anti-alpha I spectrin antibody (Abcam, Cambridge, MA) at 2×10_7_ erythrocytes/mL. Samples were washed then incubated with the fluorescently conjugated secondary antibody, goat anti-mouse Alexa Fluor 594 (Abcam, Cambridge, MA) at 1:1000 dilution. Following additional washes, samples were incubated with the fluorescently conjugated CD235a-PE antibody (Thermo Fisher Scientific, Waltham, MA) at 1:100 and SYBR Green (Thermo Fisher Scientific, Waltham, MA) at 1:10,000. A wet mount was prepared on microscope slides, and samples were immediately imaged. Images were acquired on an EVOS FL Cell Imaging System (Thermo Fisher Scientific, Waltham, MA) using RFP, GFP, and Texas Red light cubes.

### Brightfield Microscopy

Slides for brightfield imaging were prepared by fixation in methanol for 1 minute prior to the addition of Giemsa stain for 20 minutes (Sigma Aldrich, St Louis, MO). Brightfield images were obtained on an Eclipse Ci microscope (Nikon, Melville, NY) using an ImagingSource microscope camera and NIS Elements Imaging Software (Nikon, Melville, NY). For parasite stage quantification, slides were prepared for brightfield microscopy as described above both prior to and after SLOPE enrichment. Parasites were categorized by eye as either a ring, trophozoite, or schizont. In the case of a multiply infected erythrocytes, all parasites were counted separately. Gametocytes were excluded from counting.

### Re-culturing of SLOPE enriched parasites

*P. falciparum* cultures were synchronized using 5% D-sorbitol immediately prior to use. The synchronized culture was split to <1% parasitemia and 3% hematocrit to generate an untreated control flask, while the remaining 9 mL of culture were enriched according to the SLOPE protocol described above. Following enrichment, the parasite density of was measured by SYBR-Green based flow cytometry and a “SLOPE” flask was seeded with the number of the enriched cells necessary to match the parasite density of the respective untreated control flask. Uninfected erythrocytes were added to the SLOPE flask to bring hematocrit up to 3%. The parasitemia of the flasks was measured every 48h for 3 replication cycles. Untreated and SLOPE flasks were diluted every 48h to maintain parasitemia levels under 5%. For each dilution event, both flasks were subjected to the same dilution factor. The significance between groups was calculated in Graphpad Prism using a paired t-test.

### Mitochondrial membrane potential measurements

The mitochondrial membrane potential (MMP) was assessed using the MitoProbe DiIC1(5) kit (Thermo Fisher Scientific, Waltham, MA). MitoProbe DiIC1(5) accumulates in eukaryotic mitochondria in a mitochondrial membrane potential-dependent manner (53). The effect of SLOPE enrichment on MMP was tested using ring stage synchronized *P. falciparum* cultures. Following SLOPE enrichment, parasites were incubated with 50nM Mitoprobe DiIC1 (5) in media at ~1×10_6_ parasites/mL for 30 min at 37°C. Negative controls were treated with the oxidative phosphorylation uncoupler, Carbonyl cyanide m-chlorophenyl hydrazone (CCCP) (Thermo Fisher Scientific, Waltham, MA). Untreated positive controls and were assayed alongside enriched samples. Samples were co-stained with SYBR Green and analyzed using 488nm (for SYBR-Green) and 640nm (for Mitoprobe) filters on a BD Accuri C6 flow cytometer. The percentage of MMP positive parasites was determined as the ratio of Mitoprobe positive to SYBR-Green positive events. The significance between SLOPE treated and untreated groups was calculated in Graphpad Prism using a paired t-test.

### Targeted Metabolomics

*P. falciparum* parasites were synchronized using D-sorbitol 40h prior to sample collection and again, immediately prior to sample collection. Synchronized cultures at 3% parasitemia were either taken directly from culture for untreated samples or were SLOPE enriched to 45% parasitemia. 3.5×10_8_ erythrocytes were taken per sample. Samples were lysed in 0.15% saponin as previously described (54). A series of three wash steps was then performed on all samples using PBS to remove soluble erythrocyte metabolites. Metabolites were immediately extracted from the lysed pellet according to the Biocrates p180 Cell Lysis standard operating procedures. Briefly, pellets were resuspended in ice-cold ethanol/0.01 M phosphate buffer (85:15, v/v). Samples were sonicated for 3 minutes, then snap frozen in liquid nitrogen for 30 seconds. Sonication and freezing were repeated followed by a final sonication and centrifugation to pellet insoluble material. The supernatant was taken and stored at −80°C until the day of use.

Metabolite extracts were run using the AbsoluteIDQ p180 kit according to the user manual (Biocrates Life Sciences AG, Innsbruck, Austria). Samples were added to filter spots in each well along with provided internal standards and placed under a gentle stream of nitrogen gas to dry. Fresh phenyl isothiocyanate was then used for derivatization of amines, followed again by plate drying under nitrogen gas. Samples were extracted off of the filter spots using 5mM ammonium acetate in methanol. Sample extracts were either diluted 1:2 with water for LC-MS/MS analysis or diluted 1:50 with the provided flow injection analysis (FIA) mobile phase for FIA-MS/MS analysis. Liquid chromatography and mass spectrometry analysis were performed by the University of Virginia Lipidomics and Metabolomics Core. Chromatographic separation was performed using ACQUITY UPLC system (Waters Corporation, Milford, MA) with an ACQUITY 1.7 μm, 2.1 mm×75 mm BEH C18 column with an ACQUITY BEH C18 1.7 μm, 2.1mm×5mm VanGuard pre-column. Samples for both LC-MS/MS and FIA-MS/MS were analyzed on a Xevo TQ-S Mass Spectrometer (Waters Corporation, Milford, MA) according to the standard operating procedure provided by Biocrates for the AbsoluteIDQ p180kit. All metabolites were identified and quantified against the isotopically labeled internal standards. Raw data was computed in MetIDQ version Nitrogen (Biocrates Life Sciences AG, Innsbruck, Austria).

Statistical analysis and visualization of results was performed in R version 3.5.3 with tidyverse, vegan, and broom (55–57). Only metabolites detected in at least 50% of samples were analyzed (116 metabolites; **Data Set S1**). Concentration values were not normalized *post-hoc* as equal cell numbers were input for each sample. Missing values were replaced with half of the lowest detected value for each metabolite. Values were then log transformed, centered, and scaled (58). Statistically significant metabolites between SLOPE and untreated groups were determined using paired t-tests and the Benjamini and Hochberg method for multiple testing correction.

### Cholesterol manipulation and measurement

Methyl-β-cyclodextrin (mβCD) pre-saturated with cholesterol was purchased from Sigma Aldrich and dissolved in RPMI 1640 HEPES (Sigma Aldrich, St Louis, MO) to a concentration of 5mM. The solution was filtered then diluted with human serum (Valley Biomedical, Winchester, VA) to make a final “cholesterol-rich media” with 20% human serum and a final concentration of 4mM cholesterol saturated mβCD. *P. falciparum* parasites were incubated in this cholesterol-rich media for 48h. Samples were then taken for SLO lysis curve experiments as described above. Cholesterol levels in media formulations were quantified using the Amplex Red Cholesterol Assay Kit (Thermo Fisher Scientific, Waltham, MA) according to the manufacturer’s instructions. The assay was read using a SpectraMax i3x microplate reader (Molecular Devices, San Jose, CA) with Invitrogen black-walled, clear-bottom 96-well microplates (Thermo Fisher Scientific, Waltham, MA). Each replicate measurement of media cholesterol represents a separate heat inactivated batch of human plasma (Valley Biomedical, Winchester, VA) or a separate 0.5% AlbuMAX II Lipid Rich BSA preparation (Sigma Aldrich, St Louis, MO).

### Quiescent Parasite Analysis

Quiescent parasites were generated using the modified Teuscher, *et al*. protocol as described by Breglio, *et al.* (22, 30). Briefly, synchronized *P. falciparum* cultures were treated with 700nM dihydroartemisinin (Sigma Aldrich, St Louis, MO) in DMSO for 6h. Cells were washed three times with RPMI to remove drug before being resuspended in media and returned to culture conditions. Cultures were then treated with D-sorbitol every 24 hours for the following 72 hours to remove any actively growing parasites that did not enter quiescence. Immediately following the final sorbitol treatment, flasks were split into two aliquots. One aliquot was left unenriched to serve as the control, while the second aliquot was SLOPE enriched with 55 SLO units as described above. A portion of both untreated and SLOPE samples was then stained with SYBR-Green and MitoProbe DiIC1(5) as described above for flow cytometry analysis. The remainder of each untreated and SLOPE sample was stained for fluorescence microscopy as described below.

### Fluorescence Microscopy

Unenriched and SLOPE enriched samples containing quiescent parasites were stained with anti-CD235a-PE antibody, (Thermo Fisher Scientific, Waltham, MA) at 1:100 and 50nM MitoTracker Red (Thermo Fisher Scientific, Waltham, MA) at 37°C under 5% CO2 for 30 min. SYBR-Green (Thermo Fisher Scientific, Waltham, MA) was added for the last 15 min of incubation at 1:10,000. Erythrocytes were washed three times with PBS. A wet mount was immediately prepared on microscope slides, and samples were imaged using a Nikon Eclipse Ti spinning disk confocal microscope at 63X magnification using 488nm, 561nm, and 649nm lasers and the Nikon NIS Elements Software.

## Supporting information

Detailed SLOPE Protocol

Complete p180 Metabolite Data

## Acknowledgements

Special thanks to Michelle Warthan for culturing support and to the UVA Lipidomics and Metabolomics Core for targeted metabolomics run support. This project was supported with funding through the National Institute of Allergy and Infectious Disease R21AI119881 and the University of Virginia 3Cavaliers Grant (to JLG and CCM). ACB is supported by an institutional training grant (5T32GM008136-33).

## Supplemental Materials

### Supplemental Materials not included in this document

Supplemental Text 1. Detailed SLOPE Protocol (separate. docx attachment)

Supplemental Data Set 1. Complete p180 Metabolite Data (separate. csv attachment)

### All other Supplemental Materials are included below

**Supplemental Table 1.** 10 most influential metabolites for principal components 1 and 2.

| PC1 | PC2 |
| --- | --- |
| PC.aa.C34.1 | C18 |
| PC.aa.C38.5 | C18.2 |
| PC.aa.C38.6 | C18.1 |
| PC.aa.C32.2 | C16.1.OH |
| PC.aa.C36.5 | SM.C16.1 |
| PC.aa.C30.0 | lysoPC.a.C28.1 |
| PC.aa.C32.1 | C16 |
| PC.aa.C34.2 | SM.C20.2 |
| PC.aa.C36.3 | PC.aa.C28.1 |
| PC.ae.C38.0 | lysoPC.a.C24.0 |

**Supplemental Table 2.** Quantification of cholesterol from human plasma and AlbuMAX II Lipid-Rich BSA.

| Source | Average | S.E.M. | N |
| --- | --- | --- | --- |
| Human Plasma | 3300 µM | 650 µM | 4 |
| AlbuMAX II | 13.5 µM | 1.43 µM | 4 |

**Supplemental Figure 1.**
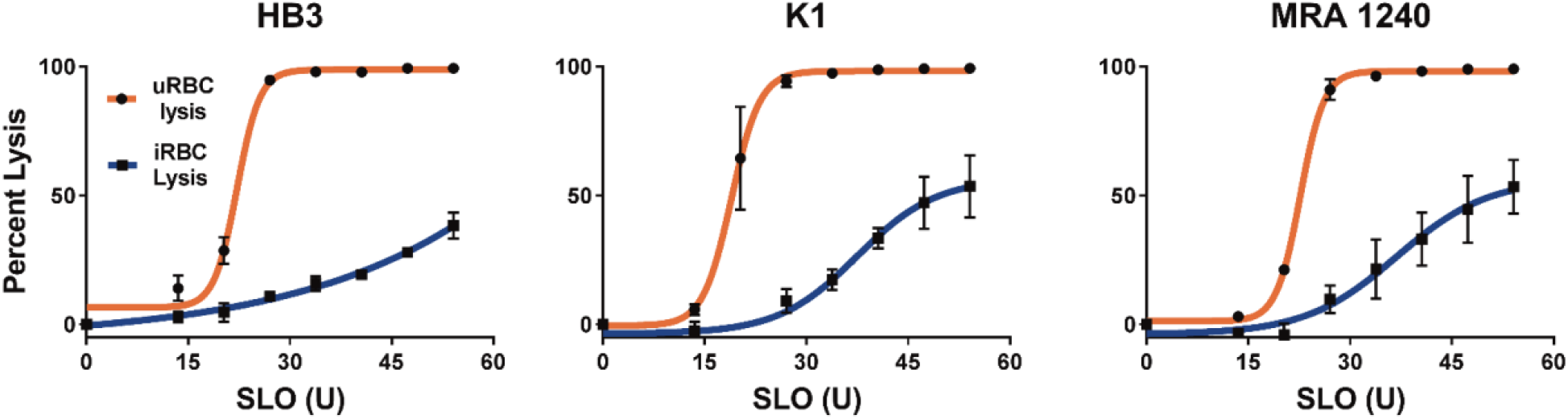
SLOPE enrichment is effective irrespective of parasite line. SLOPE enrichment performed on three different ring stage synchronized *P. falciparum* lines grown in the same blood and AlbuMAX II media batches. For each graph: *N*=3. Error bars represent S.E.M.

**Supplemental Figure 2.**
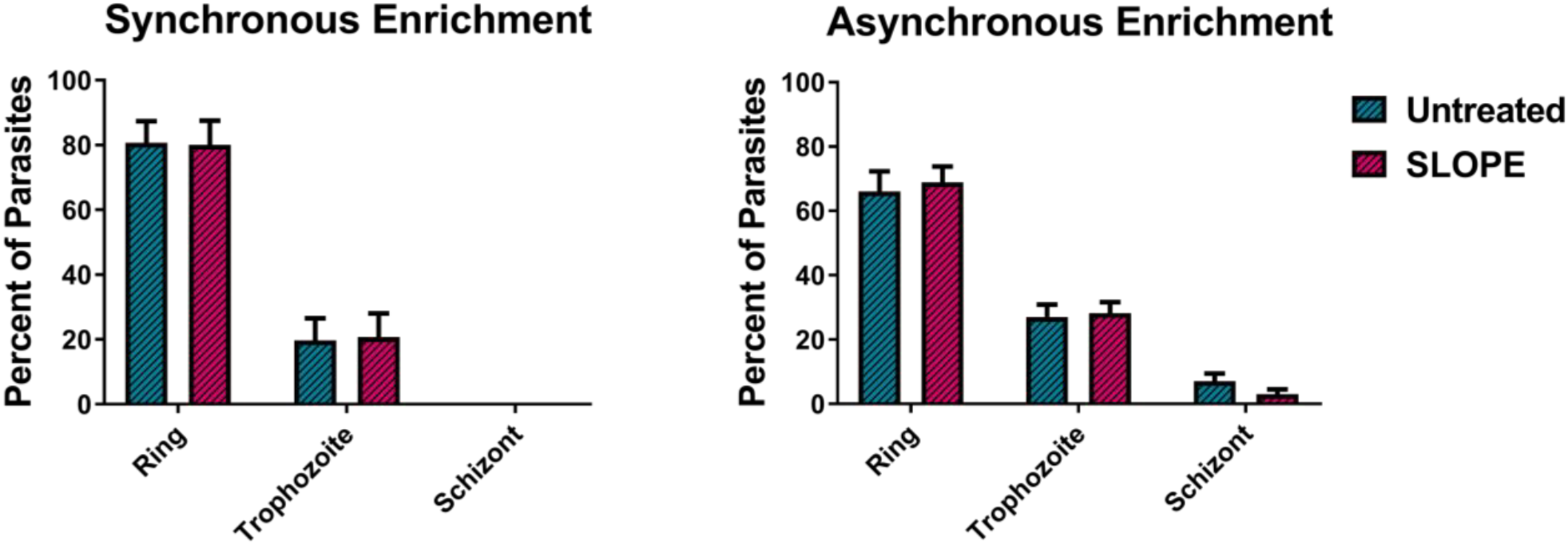
SLOPE enrichment shows no bias across stages of the intraerythrocytic cycle. *P. falciparum* line MRA 1240 cultures either (A) synchronized by one sorbitol treatment or (B) left asynchronous were staged by microscopy before and after SLOPE enrichment. For each graph: *N*=3 (200 parasites counted per replicate). Error bars represent S.E.M.

**Supplemental Figure 3.**
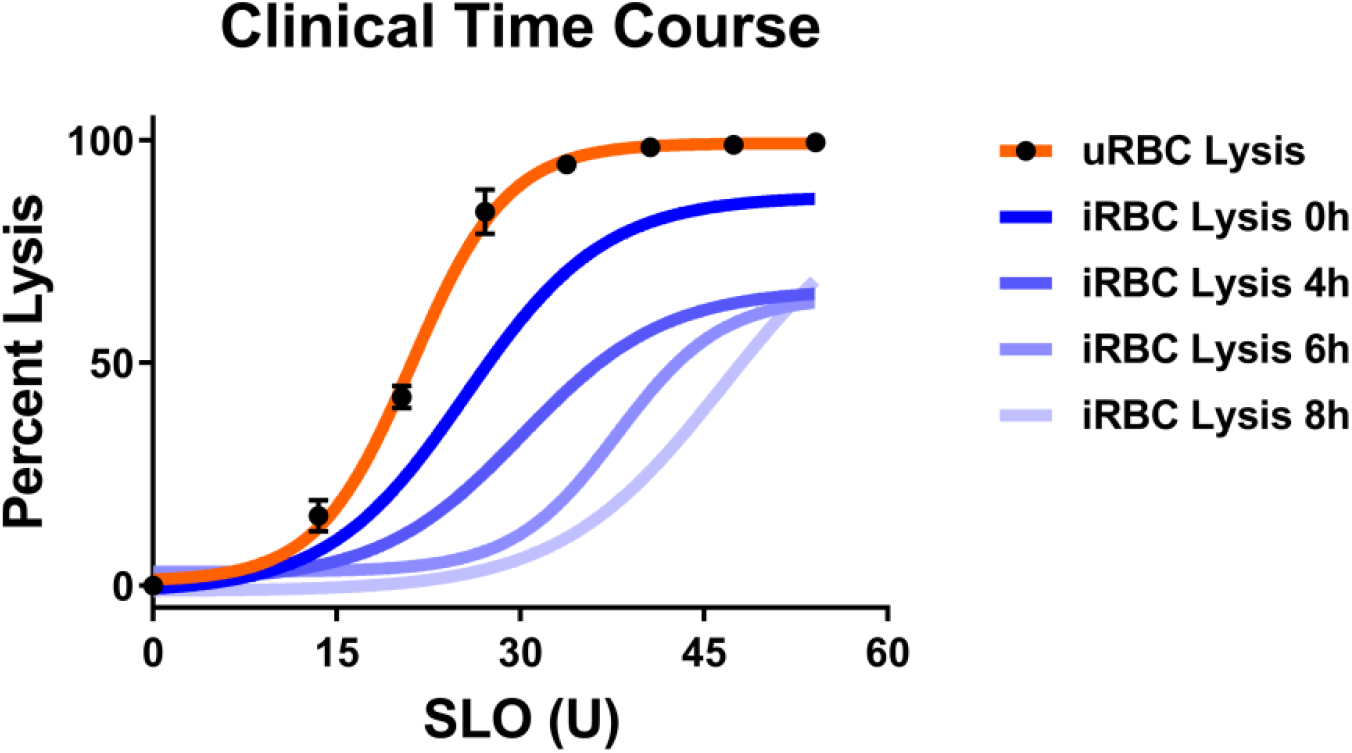
SLOPE enrichment is effective irrespective of parasite line. SLOPE enrichment performed on clinical erythroctyes incubated in cRPMI for differing lengths of time. For infected erythrocytes (iRBCs), each line represents one trial; for uninfected erythrocytes (uRBCs), lysis curves from all time points were combined as no change was observed over time: *N*=4. Error bars represent SEM.

## References

1. Organization WH. 2018. World Malaria Report 2018. WHO, Geneva, Switzerland.

2. WHO. 2015. Guidelines for the treatment of malaria. Geneva, Switzerland.

3. Blasco B, Leroy D, Fidock DA. 2017. Antimalarial drug resistance: linking Plasmodium falciparum parasite biology to the clinic. Nat Med 23:917–928.

4. Dondorp AM, Nosten F, Yi P, Das D, Phyo AP, Tarning J, Lwin KM, Ariey F, Hanpithakpong W, Lee SJ, Ringwald P, Silamut K, Imwong M, Chotivanich K, Lim P, Herdman T, An SS, Yeung S, Singhasivanon P, Day NP, Lindegardh N, Socheat D, White NJ. 2009. Artemisinin resistance in Plasmodium falciparum malaria. N Engl J Med 361:455–67.

5. Lu F, Culleton R, Zhang M, Ramaprasad A, von Seidlein L, Zhou H, Zhu G, Tang J, Liu Y, Wang W, Cao Y, Xu S, Gu Y, Li J, Zhang C, Gao Q, Menard D, Pain A, Yang H, Zhang Q, Cao J. 2017. Emergence of Indigenous Artemisinin-Resistant Plasmodium falciparum in Africa. N Engl J Med 376:991–3.

6. Prevention CfDCa. 2018. About Malaia - Biology. https://www.cdc.gov/malaria/about/biology/. Accessed

7. Ribaut C, Berry A, Chevalley S, Reybier K, Morlais I, Parzy D, Nepveu F, Benoit-Vical F, Valentin A. 2008. Concentration and purification by magnetic separation of the erythrocytic stages of all human Plasmodium species. Malar J 7:45.

8. Allman EL, Painter HJ, Samra J, Carrasquilla M, Llinas M. 2016. Metabolomic Profiling of the Malaria Box Reveals Antimalarial Target Pathways. Antimicrob Agents Chemother 60:6635–6649.

9. Cobbold SA, Chua HH, Nijagal B, Creek DJ, Ralph SA, McConville MJ. 2016. Metabolic Dysregulation Induced in Plasmodium falciparum by Dihydroartemisinin and Other Front-Line Antimalarial Drugs. J Infect Dis 213:276–86.

10. Creek DJ, Chua HH, Cobbold SA, Nijagal B, MacRae JI, Dickerman BK, Gilson PR, Ralph SA, McConville MJ. 2016. Metabolomics-Based Screening of the Malaria Box Reveals both Novel and Established Mechanisms of Action. Antimicrob Agents Chemother 60:6650–6663.

11. Olszewski KL, Morrisey JM, Wilinski D, Burns JM, Vaidya AB, Rabinowitz JD, Llinas M. 2009. Host-parasite interactions revealed by Plasmodium falciparum metabolomics. Cell Host Microbe 5:191–9.

12. Park YH, Shi YP, Liang B, Medriano CA, Jeon YH, Torres E, Uppal K, Slutsker L, Jones DP. 2015. High-resolution metabolomics to discover potential parasite-specific biomarkers in a Plasmodium falciparum erythrocytic stage culture system. Malar J 14:122.

13. Teng R, Lehane AM, Winterberg M, Shafik SH, Summers RL, Martin RE, van Schalkwyk DA, Junankar PR, Kirk K. 2014. 1H-NMR metabolite profiles of different strains of Plasmodium falciparum. Biosci Rep 34:e00150.

14. Parvazi S, Sadeghi S, Azadi M, Mohammadi M, Arjmand M, Vahabi F, Sadeghzadeh S, Zamani Z. 2016. The Effect of Aqueous Extract of Cinnamon on the Metabolome of Plasmodium falciparum Using (1)HNMR Spectroscopy. Journal of Tropical Medicine 2016:3174841.

15. Babbitt SE, Altenhofen L, Cobbold SA, Istvan ES, Fennell C, Doerig C, Llinas M, Goldberg DE. 2012. Plasmodium falciparum responds to amino acid starvation by entering into a hibernatory state. Proc Natl Acad Sci U S A 109:E3278–87.

16. Siddiqui G, Srivastava A, Russell AS, Creek DJ. 2017. Multi-omics Based Identification of Specific Biochemical Changes Associated With PfKelch13-Mutant Artemisinin-Resistant Plasmodium falciparum. J Infect Dis 215:1435–1444.

17. Carey MA, Covelli V, Brown A, Medlock GL, Haaren M, Cooper JG, Papin JA, Guler JL. 2018. Influential Parameters for the Analysis of Intracellular Parasite Metabolomics. mSphere 3.

18. Dogovski C, Xie SC, Burgio G, Bridgford J, Mok S, McCaw JM, Chotivanich K, Kenny S, Gnadig N, Straimer J, Bozdech Z, Fidock DA, Simpson JA, Dondorp AM, Foote S, Klonis N, Tilley L. 2015. Targeting the cell stress response of Plasmodium falciparum to overcome artemisinin resistance. PLoS Biol 13:e1002132.

19. David PH, Hommel M, Miller LH, Udeinya IJ, Oligino LD. 1983. Parasite sequestration in Plasmodium falciparum malaria: spleen and antibody modulation of cytoadherence of infected erythrocytes. Proceedings of the National Academy of Sciences of the United States of America 80:5075–5079.

20. Daily J. 2019. Treatment of uncomplicated falciparum malaria in nonpregnant adults and children. https://www.uptodate.com/contents/treatment-of-uncomplicated-falciparum-malaria-in-nonpregnant-adults-and-children. Accessed

21. Codd A, Teuscher F, Kyle DE, Cheng Q, Gatton ML. 2011. Artemisinin-induced parasite dormancy: a plausible mechanism for treatment failure. Malar J 10:56.

22. Teuscher F, Gatton ML, Chen N, Peters J, Kyle DE, Cheng Q. 2010. Artemisinin-induced dormancy in plasmodium falciparum: duration, recovery rates, and implications in treatment failure. J Infect Dis 202:1362–8.

23. Witkowski B, Lelievre J, Barragan MJ, Laurent V, Su XZ, Berry A, Benoit-Vical F. 2010. Increased tolerance to artemisinin in Plasmodium falciparum is mediated by a quiescence mechanism. Antimicrob Agents Chemother 54:1872–7.

24. Jackson KE, Spielmann T, Hanssen E, Adisa A, Separovic F, Dixon MW, Trenholme KR, Hawthorne PL, Gardiner DL, Gilberger T, Tilley L. 2007. Selective permeabilization of the host cell membrane of Plasmodium falciparum-infected red blood cells with streptolysin O and equinatoxin II. Biochem J 403:167–75.

25. Gulati S, Ekland EH, Ruggles KV, Chan RB, Jayabalasingham B, Zhou B, Mantel PY, Lee MC, Spottiswoode N, Coburn-Flynn O, Hjelmqvist D, Worgall TS, Marti M, Di Paolo G, Fidock DA. 2015. Profiling the Essential Nature of Lipid Metabolism in Asexual Blood and Gametocyte Stages of Plasmodium falciparum. Cell Host Microbe 18:371–81.

26. Vial HJ, Eldin P, Tielens AG, van Hellemond JJ. 2003. Phospholipids in parasitic protozoa. Mol Biochem Parasitol 126:143–54.

27. Krugliak M, Zhang J, Ginsburg H. 2002. Intraerythrocytic Plasmodium falciparum utilizes only a fraction of the amino acids derived from the digestion of host cell cytosol for the biosynthesis of its proteins. Molecular and Biochemical Parasitology 119:249–256.

28. Phillips MA. 2018. Polyamines in protozoan pathogens. Journal of Biological Chemistry 293:18746–18756.

29. Ramya TNC, Surolia N, Surolia A. 2006. Polyamine synthesis and salvage pathways in the malaria parasite Plasmodium falciparum. Biochemical and Biophysical Research Communications 348:579–584.

30. Breglio KF, Rahman RS, Sa JM, Roberts DJ, Wellems TE. 2018. Kelch Mutations in Plasmodium falciparum Protein K13 Do Not Modulate Dormancy after Artemisinin Exposure and Sorbitol Selection In Vitro. Antimicrob Agents Chemother 62.

31. Gray KA, Gresty KJ, Chen N, Zhang V, Gutteridge CE, Peatey CL, Chavchich M, Waters NC, Cheng Q. 2016. Correlation between Cyclin Dependent Kinases and Artemisinin-Induced Dormancy in Plasmodium falciparum In Vitro. PLoS One 11:e0157906.

32. Peatey CL, Chavchich M, Chen N, Gresty KJ, Gray KA, Gatton ML, Waters NC, Cheng Q. 2015. Mitochondrial Membrane Potential in a Small Subset of Artemisinin-Induced Dormant Plasmodium falciparum Parasites In Vitro. J Infect Dis 212:426–34.

33. Tokumasu F, Crivat G, Ackerman H, Hwang J, Wellems TE. 2014. Inward cholesterol gradient of the membrane system in P. falciparum-infected erythrocytes involves a dilution effect from parasite-produced lipids. Biol Open 3:529–41.

34. Quarfordt SH, Hilderman HL. 1970. Quantitation of the in vitro free cholesterol exchange of human red cells and lipoproteins. J Lipid Res 11:528–35.

35. Mbengue A, Bhattacharjee S, Pandharkar T, Liu H, Estiu G, Stahelin RV, Rizk SS, Njimoh DL, Ryan Y, Chotivanich K, Nguon C, Ghorbal M, Lopez-Rubio JJ, Pfrender M, Emrich S, Mohandas N, Dondorp AM, Wiest O, Haldar K. 2015. A molecular mechanism of artemisinin resistance in Plasmodium falciparum malaria. Nature 520:683–7.

36. Witkowski B, Amaratunga C, Khim N, Sreng S, Chim P, Kim S, Lim P, Mao S, Sopha C, Sam B, Anderson JM, Duong S, Chuor CM, Taylor WRJ, Suon S, Mercereau-Puijalon O, Fairhurst RM, Menard D. 2013. Novel phenotypic assays for the detection of artemisinin-resistant Plasmodium falciparum malaria in Cambodia: in-vitro and ex-vivo drug-response studies. The Lancet Infectious diseases 13:1043–1049.

37. Claessens A, Affara M, Assefa SA, Kwiatkowski DP, Conway DJ. 2017. Culture adaptation of malaria parasites selects for convergent loss-of-function mutants. Scientific reports 7:41303–41303.

38. Daily JP, Scanfeld D, Pochet N, Le Roch K, Plouffe D, Kamal M, Sarr O, Mboup S, Ndir O, Wypij D, Levasseur K, Thomas E, Tamayo P, Dong C, Zhou Y, Lander ES, Ndiaye D, Wirth D, Winzeler EA, Mesirov JP, Regev A. 2007. Distinct physiological states of Plasmodium falciparum in malaria-infected patients. Nature 450:1091.

39. Peters JM, Fowler EV, Krause DR, Cheng Q, Gatton ML. 2007. Differential Changes in Plasmodium falciparum var Transcription during Adaptation to Culture. The Journal of Infectious Diseases 195:748–755.

40. White J, Mascarenhas A, Pereira L, Dash R, Walke JT, Gawas P, Sharma A, Manoharan SK, Guler JL, Maki JN, Kumar A, Mahanta J, Valecha N, Dubhashi N, Vaz M, Gomes E, Chery L, Rathod PK. 2016. In vitro adaptation of Plasmodium falciparum reveal variations in cultivability. Malaria Journal 15:33.

41. Barrett MP, Kyle DE, Sibley LD, Radke JB, Tarleton RL. 2019. Protozoan persister-like cells and drug treatment failure. Nature Reviews Microbiology doi:10.1038/s41579-019-0238-x.

42. Chen N, LaCrue AN, Teuscher F, Waters NC, Gatton ML, Kyle DE, Cheng Q. 2014. Fatty acid synthesis and pyruvate metabolism pathways remain active in dihydroartemisinin-induced dormant ring stages of Plasmodium falciparum. Antimicrob Agents Chemother 58:4773–81.

43. Cooper KD, Shukla JB, Rennert OM. 1976. Polyamine distribution in cellular compartments of blood and in aging erythrocytes. Clin Chim Acta 73:71–88.

44. Shen H-M, Chen S-B, Cui Y-B, Xu B, Kassegne K, Abe EM, Wang Y, Chen J-H. 2018. Whole-genome sequencing and analysis of Plasmodium falciparum isolates from China-Myanmar border area. Infectious diseases of poverty 7:118–118.

45. Coppens I, Joiner KA. 2003. Host but not parasite cholesterol controls Toxoplasma cell entry by modulating organelle discharge. Molecular biology of the cell 14:3804–3820.

46. Hirsch JG, Bernheimer AW, Weissmann G. 1963. MOTION PICTURE STUDY OF THE TOXIC ACTION OF STREPTOLYSINS ON LEUCOCYTES. J Exp Med 118:223–8.

47. Launay J-M, Alouf JE. 1979. Biochemical and ultrastructural study of the disruption of blood platelets by streptolysin O. Biochimica et Biophysica Acta (BBA) - Biomembranes 556:278–291.

48. Lim C, Hansen E, DeSimone TM, Moreno Y, Junker K, Bei A, Brugnara C, Buckee CO, Duraisingh MT. 2013. Expansion of host cellular niche can drive adaptation of a zoonotic malaria parasite to humans. Nature Communications 4:1638.

49. Trager W, Jensen JB. 1976. Human malaria parasites in continuous culture. Science 193:673–5.

50. Bei AK, Desimone TM, Badiane AS, Ahouidi AD, Dieye T, Ndiaye D, Sarr O, Ndir O, Mboup S, Duraisingh MT. 2010. A flow cytometry-based assay for measuring invasion of red blood cells by Plasmodium falciparum. Am J Hematol 85:234–7.

51. Kassaza K, Operario DJ, Nyehangane D, Coffey KC, Namugosa M, Turkheimer L, Ojuka P, Orikiriza P, Mwanga-Amumpaire J, Byarugaba F, Bazira J, Guler JL, Moore CC, Boum Y. 2018. Detection of *Plasmodium* Species by High-Resolution Melt Analysis of DNA from Blood Smears Acquired in Southwestern Uganda. Journal of Clinical Microbiology 56:e01060–17.

52. Lambros C, Vanderberg JP. 1979. Synchronization of Plasmodium falciparum erythrocytic stages in culture. J Parasitol 65:418–20.

53. Grimberg BT, Jaworska MM, Hough LB, Zimmerman PA, Phillips JG. 2009. Addressing the malaria drug resistance challenge using flow cytometry to discover new antimalarials. Bioorganic & medicinal chemistry letters 19:5452–5457.

54. Doolan DL. 2002. Malaria methods and protocols, vol 72. Springer Science & Business Media.

55. David Robinson AH, Matthieu Gomez, Boris Demeshev, Dieter Menne, Benjamin Nutter, Luke Johnston, Ben Bolker, Francois Briatte, Jeffrey Arnold, Jonah Gabry, Luciano Selzer, Gavin Simpson, Jens Preussner, Jay Hesselberth, Hadley Wickham, Matthew Lincoln, Alessandro Gasparini, Lukasz Komsta, Frederick Novometsky, Wilson Freitas, Michelle Evans, Jason Cory Brunson, Simon Jackson, Ben Whalley, Michael Kuehn, Jorge Cimentada, Erle Holgersen, Karl Dunkle Werner. 2019. broom: Convert Statistical Analysis Objects into Tidy Tibbles. https://cran.r-project.org/web/packages/broom/. Accessed

56. Jari Oksanen FGB, Michael Friendly, Roeland Kindt, Pierre Legendre, Dan McGlinn, Peter R. Minchin, R. B. O’Hara, Gavin L. Simpson, Peter Solymos, M. Henry H. Stevens, Eduard Szoecs, Helene Wagner. 2019. vegan: Community Ecology Package. https://cran.r-project.org/web/packages/vegan/. Accessed

57. Wickman H. 2017. tidyverse: Easily Install and Load the ‘Tidyverse’. https://cran.r-project.org/web/packages/tidyverse/. Accessed

58. Sugimoto M, Kawakami M, Robert M, Soga T, Tomita M. 2012. Bioinformatics Tools for Mass Spectroscopy-Based Metabolomic Data Processing and Analysis. Curr Bioinform 7:96–108.

