## Supplementary material for "Cholesterol-dependent enrichment of understudied erythrocytic stages of human *Plasmodium* parasites": Detailed SLOPE Protocol

**Text S1. Detailed SLOPE Protocol.**

**Protocol Developed and Written by Audrey Brown**

**PI: Jennifer Guler, University of Virginia**

**SLO activation**

Once a new vial of SLO has been received from a vendor (ex. Sigma-Aldrich S5265), the lyophilized powder should be activated according the following protocol (adapted from Jackson, *et al*. *Biochem J.* 2007).

1. Make activation solution: 1X PBS + 0.1% BSA + 5mM DTT
2. Resuspend lyophilized powder in 5ml of activation solution. Mix well by pipetting or inverting.
3. Incubate solution at 37C for 2 hours.
4. Mix the solution well by pipetting or inverting.
5. Divide into aliquots and store at -20C.

**Defining a Hemolytic Unit (U)**

Testing to determine the SLO solution volume which constitutes a U should be done in triplicate every time a new shipment of SLO has been received and activated.

1. Preincubate uRBCs in growth media at 37C for at least 4h. (This allows uRBC cholesterol levels to accurately reflect the cholesterol levels that will occur in *in vitro* culture and accounts for cholesterol level variation between media batches).
2. Suspend uRBCs at 2% hematocrit in 1X PBS.
3. Label a series of eppendorfs with the given SLO volumes you wish to test. (We recommend testing 6-8 points between 0µl SLO and 1.5µl SLO.
4. Aliquot 50µl of 2% uRBCs into each tube.
5. Make the appropriate dilutions of SLO in 1X PBS so that equal volumes can be added to each tube. (Also add the equivalent volume of PBS to the “0µl SLO” control tube).
6. Mix each tube by pipetting.
7. Incubate at 37C for 30min.
8. Remove tubes from heat and quickly mix by pipetting.
9. Add 10µl from each tube into 990µl of 1X PBS to generate a 1:100 dilution. Dilution aids in stopping lysis. (We suggest preparing tubes for dilutions during the 30min incubation period so that this step can proceed immediately following the end of the incubation).
10. Use the Cellometer T4 to count the intact RBC cell density of each sample.

*If a Cellometer T4 is not available, intact cells can be counted by flow cytometry. Alternatively, lysis can be measured spectrophotometrically by centrifuging (undiluted) tubes and taking aliquots of supernatant. Then measure samples at 412nm to determine hemoglobin release. Finally, normalize against a sample subjected to hypo-osmotic lysis (see Jackson, *et al.* *Biochem J*. 2007).

11. Determine the fraction of cells lysed by subtracting the cell density of each SLO-containing sample from the cell density of the “0µl SLO” control tube. Then divide this value by the “0µl SLO” control cell density.
12. Plot cell lysis as a function of SLO amount to determine the amount of SLO which causes 50% lysis. This is 1 hemolytic unit.

**SLOPE enrichment**

1. Count the cell density of your sample using a flow cytometer, hemocytometer, or Cellometer T4.

***Note**: SLOPE enrichment will uniformly enrich for all asexual blood stages. If you wish to enrich a specific stage, such as only ring stage parasites, utilize your synchronization method of choice prior to this step.

2. Spin down the sample and remove cell culture media. Add the volume of non-cholesterol containing solution (ex. PBS, RPMI) required to yield a concentration of 2x10^9^ RBCs/ml.
3. For every 5 volumes of RBCs you wish to SLOPE enrich, SLO and 1X PBS should be added to a total of 2 volumes.
   a. Ex. Given an SLO stock of 2U/µl, a desired SLO activity of 30U, and cell amount of 1x10^8^ RBCs for enrichment: to 50µl RBCs (at 2x10^9^ RBCs/ml), add 15µl SLO (30U) and 5µl PBS (SLO + PBS = 20µl total aka “2 volumes”).
4. Mix by pipetting and incubate at room temperature for exactly 6 min.
5. Add >5-10 volumes of PBS or RPMI and mix by pipetting
6. Spin at 2,500xg for ~3 min, remove supernatant

*Delaying the initial wash may lead to over lysis.

7. Add PBS or RPMI to wash, spin at 2,500xg for 3 min, remove supernatant.
8. Repeat step 7 for a total of 3 washes.

*The wash step can be repeated again if lysis appears to still be occurring as evidenced by hemoglobin in the supernatant. (Even though the free SLO has been removed, lysis may continue to take place for a few minutes). Ensure lysis has finished before proceeding to the Percoll gradient step.

9. Resuspend cells in desired volume of PBS or media (cholesterol containing solutions, such as serum-supplemented RPMI, may now be used).
10. Slowly layer cells on top of a 60% Percoll gradient.
    a. To make Percoll: take 9 parts Percoll and add 1-part 10X PBS to make stock isotonic Percoll (SIP).
    b. Dilute from SIP to make 60% Percoll using 1X PBS (2-parts SIP plus 1-part 1X PBS)
11. Spin the Percoll gradient at 1500xg for 5min using a swinging bucket rotor. (Spin may need to be extended for >5min, if you are using a large Percoll gradient, ex. 50ml).
12. Carefully remove and discard the upper fraction of Percoll, which contains lysed RBC ghosts. Keep the pipette tip at the top of the Percoll as you are removing this fraction to ensure ghosts are removed effectively.
13. Using a new pipette tip to avoid cross-contamination of ghosts, remove the lower, intact RBC fraction and place these cells into a clean tube.
14. Wash cells twice in >10 volumes of PBS or media.
15. Optional: Take a small aliquot to assess parasitemia by Giemsa staining a blood smear or by SYBR Green-based flow cytometry.

*Parasites can now be used for downstream applications.
